# DNA damage-associated protein co-expression network in cardiomyocytes informs on tolerance to genetic variation and disease

**DOI:** 10.1101/2024.08.14.607863

**Authors:** Omar D. Johnson, Sayan Paul, Jose A. Gutierrez, William K. Russell, Michelle C. Ward

**Affiliations:** Biochemistry, Cellular and Molecular Biology Graduate Program, University of Texas Medical Branch, Galveston, Texas, USA; MD-PhD Combined Degree Program, University of Texas Medical Branch, Galveston, Texas, USA; Department of Biochemistry and Molecular Biology, University of Texas Medical Branch, Galveston, Texas, USA

**Author notes:** Correspondance should be addressed to M.C.W. These authors contributed equally to the work (Omar D. Johnson, Sayan Paul).

## Abstract

Cardiovascular disease (CVD) is associated with both genetic variants and environmental factors. One unifying consequence of the molecular risk factors in CVD is DNA damage, which must be repaired by DNA damage response proteins. However, the impact of DNA damage on global cardiomyocyte protein abundance, and its relationship to CVD risk remains unclear. We therefore treated induced pluripotent stem cell-derived cardiomyocytes with the DNA-damaging agent Doxorubicin (DOX) and a vehicle control, and identified 4,178 proteins that contribute to a network comprising 12 co-expressed modules and 403 hub proteins with high intramodular connectivity. Five modules correlate with DOX and represent distinct biological processes including RNA processing, chromatin regulation and metabolism. DOX-correlated hub proteins are depleted for proteins that vary in expression across individuals due to genetic variation but are enriched for proteins encoded by loss-of-function intolerant genes. While proteins associated with genetic risk for CVD, such as arrhythmia are enriched in specific DOX-correlated modules, DOX-correlated hub proteins are not enriched for known CVD risk proteins. Instead, they are enriched among proteins that physically interact with CVD risk proteins. Our data demonstrate that DNA damage in cardiomyocytes induces diverse effects on biological processes through protein co-expression modules that are relevant for CVD, and that the level of protein connectivity in DNA damage-associated modules influences the tolerance to genetic variation.

## Introduction

Cardiovascular disease (CVD) is the leading cause of mortality globally (1). CVD risk factors including age, diagnosis of metabolic disease, and treatment with chemotherapeutics can result in oxidative stress and apoptosis leading to accumulated DNA damage and activation of the DNA damage response in the heart (2). The amount of DNA damage in the myocardium, as determined by the DNA double-strand break (DSB) marker gamma γH2AX, is predictive of heart failure (3). Persistent DNA damage in cardiac cell types can lead to disease and has been detected among some of the most prevalent CVDs, including heart failure, dilated cardiomyopathy, and atrial fibrillation (4). The myocardium is comprised of cardiomyocytes that produce the contractile force of the heart necessary to circulate oxygenated blood throughout the body, and therefore damage to these cells can lead to cardiac dysfunction. Adult human cardiomyocytes are also particularly susceptible to DNA damage given that they are post-mitotic and unable to regenerate (4). This means that DNA damage, induced through DSBs, can only be repaired through error-prone Non-homologous end joining, unlike proliferative cell types that can also repair DNA through homologous recombination (5).

Doxorubicin (DOX) is an effective anthracycline chemotherapeutic that can adversely induce cardiac dysfunction through the formation of DSBs in cardiomyocytes (6). This is primarily mediated through its interaction with the DNA topology regulator topoisomerase II (TOP2). Physiologically, the TOP2A and TOP2B isoforms resolve torsional stress in DNA by DSBs; however, in the presence of DOX, TOP2 is trapped on DNA where it generates DNA lesions (7). The predominant TOP2 isoform expressed in the heart is TOP2B. TOP2B has been shown to mediate the cardiotoxic effects of DOX in *in vivo* animal models and *in vitro* human disease models (8, 9). While DOX can lead to cellular effects through mechanisms including the generation of ROS, at clinically-tolerated sub-micromolar doses, DSB induced through interactions with TOP2B is the main contributor to DOX-induced cardiotoxicity (10). Clinically, DOX-induced cardiac dysfunction shares characteristics of multiple CVDs (6). Nine percent of individuals receiving DOX exhibit reductions in their left ventricular ejection fractions within values that would constitute heart failure (11, 12). Similarly, treatment with DOX increases the risk for electrophysiologic dysfunction and atrial fibrillation by tenfold, and is associated with other clinically-measurable phenotypes, such as an increased QT-interval (13, 14). These pathologies overlap with those influenced by DNA damage and are impacted by genetic risk (4, 15).

Genome-wide association studies (GWAS) have identified hundreds of risk loci associated with complex CVDs including atrial fibrillation, heart failure, DOX-induced cardiotoxicity and clinical cardiovascular phenotypes, highlighting the genetic component of CVD (15). Although GWAS identify genetic risk loci that can be mapped to genes that implicate putative regulatory effects, they do not explain the molecular mechanisms of the diseases that they associate with (16, 17). This ultimately impedes our understanding of the effect of genetic variation on CVD and its applicability to identify potential drug targets (17).

One approach to understand a complex disease phenotype is to construct networks based on molecular phenotypes such as global mRNA or protein expression levels. Indeed, transcriptome profiling of human brain regions has provided insight into cell type specificity for the risk for neuropsychiatric diseases (18). However, it has been shown that expression networks differ at the mRNA and protein level and that many complex disease phenotypes are observed only at the proteome level (19–21). Targeted studies investigating the protein interactomes of proteins encoded in GWAS loci have identified convergent points in the interaction network for autism spectrum disorder and coronary artery disease highlighting disease-relevant biology (22, 23). Complex disease networks are likely to exert tissue and cell type-specific effects. Indeed, proteins relevant to late onset Alzheimer’s disease are localized within glial cells in the brain (21). Similarly, the appropriate cellular context, such as cell type and state, is important for understanding the basis of CVD. Clinically-observed CVD such as DOX-induced cardiotoxicity can be recapitulated in human induced pluripotent stem cell-derived cardiomyocyte models (24), allowing for the study of disease-relevant states such as exposure to DOX and hypoxia to understand CVD risk (25–27). However, a protein network has not been generated in this context to understand the interplay between a cellular stressor relevant to CVD and proteins encoded by genes that are implicated in complex CVDs.

We therefore designed a study to determine the effects of the DNA-damaging drug, DOX, on the proteome of human cardiomyocytes. We differentiated cardiomyocytes from induced pluripotent stem cells from three healthy individuals, treated them with a sub-lethal dose of DOX, and measured global protein expression levels. We were able to construct a protein expression network consisting of co-expression modules correlated with DNA damage treatment that associate with distinct biological processes and cardiovascular traits and diseases, and define the tolerance of network components to genetic variation.

## Results

### iPSC-CM proteome resembles the heart ventricle proteome and is affected by DNA damage

We differentiated induced pluripotent stem cells (iPSCs) from three healthy female donors into cardiomyocytes (iPSC-CMs) using biphasic WNT modulation (Fig 1A, Table 1). iPSC-CMs were metabolically selected and matured for 27 days post-differentiation initiation (See Methods) (26). Flow cytometry analysis of two individuals indicated high-purity cultures with a median of 97% of cells expressing cardiac troponin T (Fig S1).

**Figure 1:**
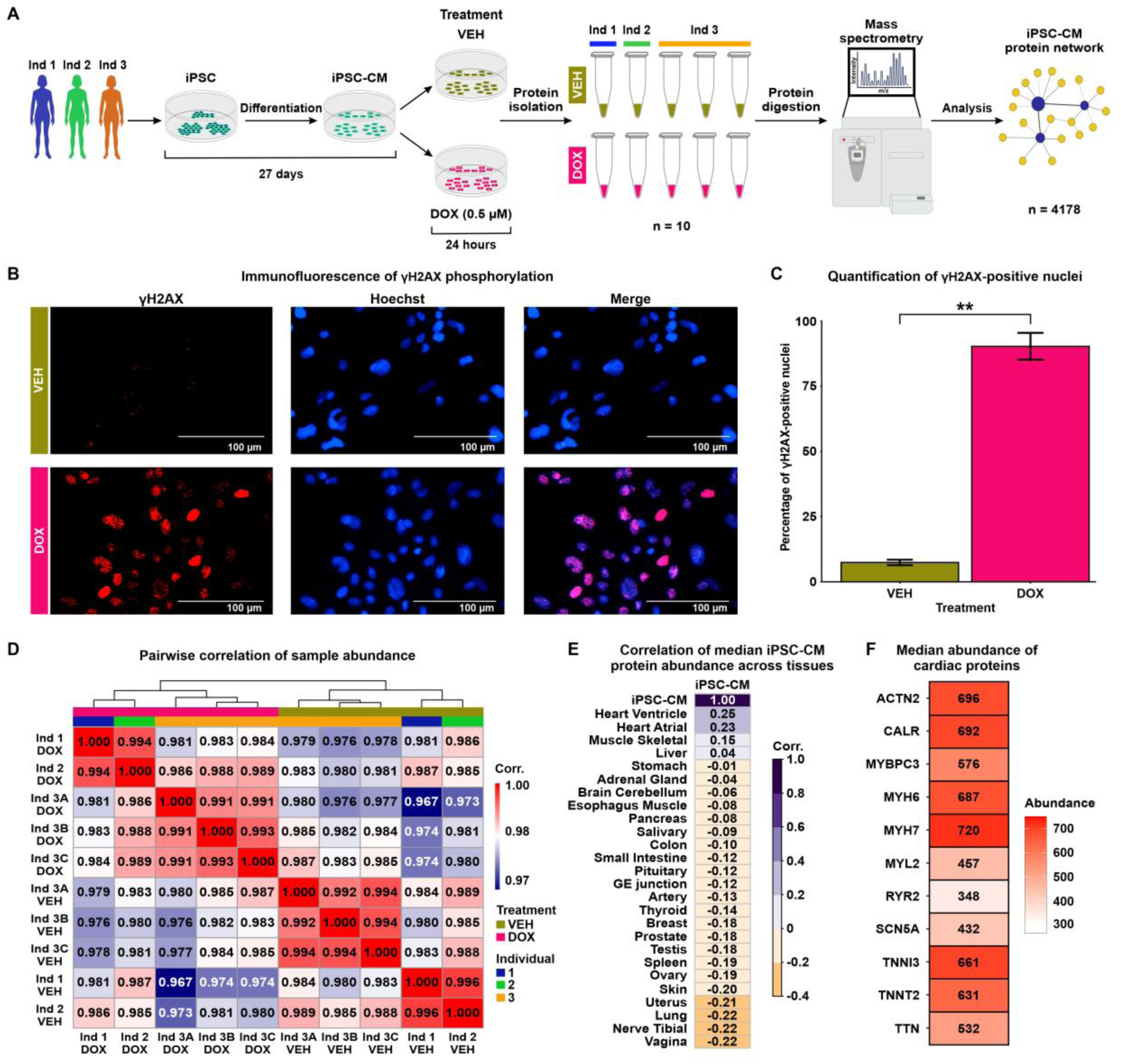
iPSC-CM protein samples cluster by DOX treatment and most closely resemble the heart ventricle proteome. **(A)** Flowchart representing the study design. iPSCs from three individuals (Ind) Ind 1 (blue), Ind 2 (green) and Ind 3 (orange) were differentiated into cardiomyocytes (iPSC-CMs). iPSC-CMs were exposed to 0.5 μM Doxorubicin (DOX) or a vehicle control (VEH) for 24 hours. The treatment was replicated in Ind 3 three times, yielding 10 total samples. Peptides were isolated and quantified by mass spectrometry allowing the construction of an iPSC-CM network from 4,178 proteins. **(B)** Immunostaining of the DNA damage marker, and Hoechst nuclear stain in VEH- and DOX-treated iPSC-CMs. **(C)** Percentage of VEH- and DOX-treated iPSC-CMs that stain positive for γH2AX. Data representative of treatment experiments from three individuals. Asterisk represents a statistically significant change in γH2AX expression (\*\**P* < 0.01). **(D)** Pairwise Pearson correlation of the median protein abundance across all 10 samples. **(E)** Pearson correlation of the median iPSC-CM protein abundance for all proteins across all experimental samples to the median abundance of those proteins across different human postmortem tissues (20). **(F)** Median protein abundance across experimental samples of select proteins known to be elevated in heart tissue in comparison to other tissue types.

To determine the effect of DNA damage on the cardiomyocyte proteome, we treated iPSC-CMs with a TOP2B-inhibiting, clinically-relevant concentration of DOX (0.5 μM) and a water vehicle control (VEH) for 24 hours. This dose of DOX causes minimal cell death in iPSC-CMs but induces thousands of mRNA expression changes for genes in pathways related to p53 signaling, base excision repair and DNA replication (28). We confirmed the DNA-damaging effects of DOX under these conditions by assaying the expression of the DNA DSB marker γH2AX (Fig 1B). DOX-treated cardiomyocytes have significantly higher γH2AX expression compared to VEH-treated cardiomyocytes (Fig 1C; 90% vs. 7%; t-test; *P* < 0.05). To account for technical variability in the drug treatment and proteomic data collection, the DOX and VEH treatment in iPSC-CMs from one individual was replicated three times, resulting in a total of 10 samples across individuals and treatments.

Global protein expression data was inferred from peptidic identification and quantification using data-independent acquisition mass spectrometry (DIA; See Methods). Peptides were mapped to 4,261 proteins present in at least one sample (S1 Appendix). Four non-human proteins and proteins that were present in less than half of the samples were filtered out. To enable construction of a complete as possible network, we used the remaining 4,178 proteins to impute protein abundance data for the 246 proteins with missing data using a feature clustering-based imputation method commonly used for proteomics data (29, 30) (Fig S2A). On average, imputed proteins are less abundant than proteins present in all 10 samples (median log_2_ abundance all = 19.83, median imputed abundance = 16.98; Fig S2B). We took advantage of the technical replicates to remove unwanted variation in the data (See Methods) (31). After correction, principal component analysis reveals that PC1, which accounts for 32% of variation in the data, associates with drug treatment, while PC2, accounting for 29% of the variation in the data, associates with individual (Fig S2C-D). Similarly, when comparing all pairwise sample correlations, the data primarily separates into two clusters corresponding to DOX and VEH treatment (Fig 1D).

To gain insight into the utility of our iPSC-CM proteome data for understanding the effect of DNA damage on the heart, we correlated the median expression of our set of proteins with the expression of proteins measured in 26 post-mortem human tissues from hundreds of individuals (20). The iPSC-CM proteome is most similar to the proteome from heart left ventricle and atrial appendage, followed by skeletal muscle (Fig 1E). iPSC-CMs express cardiac-specific and cardiac-abundant proteins, including Myosin Heavy Chain 7 (MYH7), Myosin Light Chain 6 (MYH6), and Troponin I (TNNI3; Fig 1F). Together, these findings affirm that the proteome of our iPSC-CMs closely resembles ventricular and atrial tissue, includes key cardiomyocyte proteins, and demonstrates the influence of DNA damage as the main contributor to variation of protein abundance values within our experimental model.

### Network analysis reveals DOX-correlated modules, response proteins and hub proteins

In order to identify sets of co-expressed proteins within our data, we utilized Weighted Correlation Network Analysis (WGCNA) (32). WGCNA assumes the co-expression network follows a scale-free topology, where few nodes (proteins) have high connectivity and many nodes have low connectivity, and requires a soft power threshold to determine the weights of edges connecting nodes. We regressed total network connectivity on the connectivity frequency distribution at different power thresholds and determined that the lowest power threshold that yielded the best fit to a scale-free network (0.71 regression coefficient) was 20 (Fig S3A). The scale-free fit is further supported by decreasing connectivities at higher power thresholds, where we find that at a power threshold of 20, our network has a median connectivity score of 53.6, mean connectivity of 65.4, and max connectivity of 216 (Fig S3B). Using these criteria, we generated a network comprising 21 co-expressed modules with at least 40 proteins per module (Fig S3C), summarized by 21 eigenproteins (Fig S3D). All combinations of proteins in the co-expression network are summarized as pairwise correlations (S2 Appendix). To better distinguish individual modules, we merged modules with highly correlated eigenproteins (Pearson correlation > 0.85), which collapsed the data into 12 distinct modules (Fig 2A & Fig S3E). These modules contain between 45 and 798 proteins (Table 2). 106 proteins do not fit the profile of any module and were categorized as ‘unassigned’ (Table 2). Imputed proteins are distributed across modules representing 1.6-16% of all proteins within a module. We associated each module with the two biological features of our data, namely DOX treatment and Individual (IND). To do so, we measured the correlation between each module’s eigenprotein and either IND or DOX treatment (Fig 2A). We named each of the 12 modules based on the order of their absolute correlation with DOX treatment α, β, γ, δ, ε, ζ, η, θ, ι, κ, λ, and μ. Eigenproteins from the α, β, γ, ε, and δ modules exhibited significant correlations with DOX treatment (Pearson correlation, *P* < 0.01), and were therefore categorized as DOX-correlated modules. Modules α, δ and ε have strong negative DOX correlations of -0.97, -0.81 and -0.79, while modules β and γ have similar positive correlations of 0.89 and 0.86, respectively. In contrast, eigenproteins from the μ, ι, and θ modules are significantly correlated with IND. This delineation indicates that five of the 12 co-expressed modules in our iPSC-CM protein abundance correlation network are specifically associated with DOX treatment.

**Figure 2:**
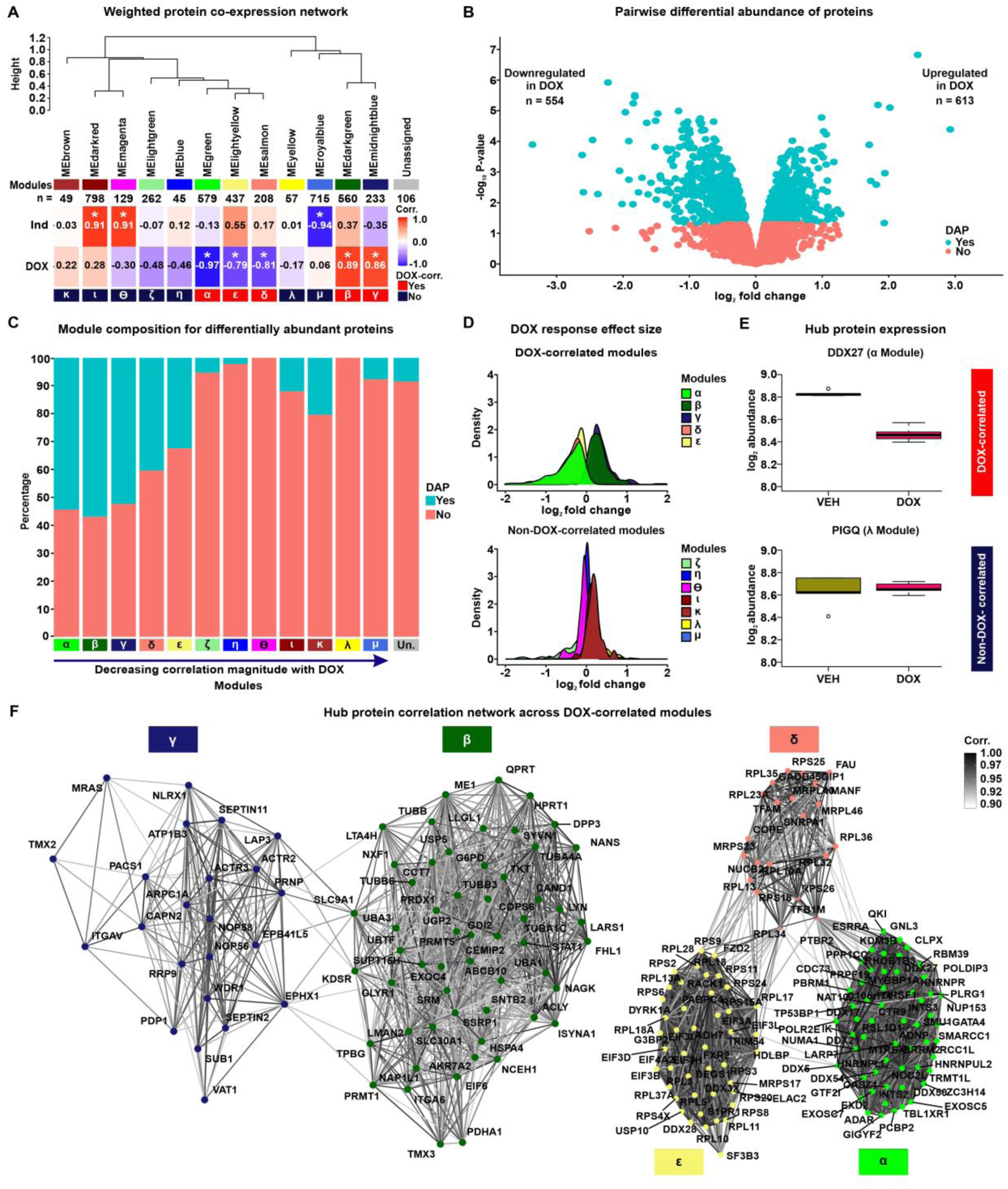
Network analysis of the iPSC-CM proteome identifies protein co-expression modules correlated with DOX treatment. **(A)** Hierarchical clustering of 12 co-expression module eigen (ME) proteins based on their Pearson correlation. Height represents the dissimilarity between ME proteins (1-corr.). Each module is represented by a color and the number of proteins in the module specified. The “Unassigned” module, shaded grey, includes proteins that cannot be represented by one of the 12 ME proteins. The correlation between each ME protein and the known biological variables: Individual (Ind) and DOX treatment is shown. Asterisk represents a significant correlation between the ME protein and the trait (\**P* < 0.01). Modules are designated by Greek letters in order of decreasing correlation with DOX and summarized as DOX-correlated modules (red), and non-DOX-correlated modules (dark blue). **(B)** Volcano plot representing proteins that are differentially abundant (DAPs; *P* < 0.05; blue) and not differentially abundant (salmon) between VEH- and DOX-treated iPSC-CMs. **(C)** Percentage of DAPs (blue) and non-DAPs (salmon) across co-expression modules where modules are ordered by decreasing correlation to DOX treatment, and amongst a module-unassigned set (grey). **(D)** Distribution of effect sizes of response to DOX treatment (log_2_ fold change from pairwise differential abundance model) for the five DOX-correlated and seven non-DOX-correlated modules. **(E)** Examples of hub protein abundance values in VEH- and DOX-treated iPSC-CMs in a DOX-correlated module (α; DDX27) and a non-DOX-correlated module (λ; PIGQ). **(F)** DOX-correlated hub protein co-expression correlation network, where nodes are hub proteins and edges represent the weighted correlation between them. Connections among DOX-correlated proteins with a correlation of ≥ 0.9 are depicted for visualization.

To support the categorization of five modules as DOX-correlated, we performed an independent pairwise differential abundance (DA) test. Using this approach, we identified 1,167 DA proteins among the 4,178 evaluated proteins (*P* < 0.05; Table 2; See Methods). 613 DA proteins exhibit increased abundance in the DOX-treated iPSC-CMs, while 554 DA proteins show decreased abundance (Fig 2B). We overlapped the set of DA proteins with the set of proteins in each module and found 33 - 57% of α, β, γ, ε, and δ DOX-correlated module proteins are classified as DA compared to 0 - 20% of non-DOX-correlated module proteins (Fig 2C). Similarly, proteins within DOX-correlated modules tend to have a greater response to DOX treatment as measured by their log fold change, compared to proteins in non-DOX-correlated modules (Fig 2D).

In order to test the robustness of our approach for identifying DOX-responsive proteins, we also acquired protein measurements of the same samples by data-dependent acquisition (DDA) on the mass spectrometer (See Methods). We identified 4,501 proteins using this method. All proteins identified by DIA were also identified by DDA, and the abundance of the 3,027 proteins present across all samples in both data sets is highly correlated (rho = 0.75, *P* < 0.001; Fig S4). Following imputation and unwanted variance removal of the DDA data, we identified 620 DA proteins amongst 3,954 proteins. The effect size of the response to DOX treatment amongst proteins included in both acquisition methods is correlated (Pearson correlation coefficient = 0.35; *P* < 0.001; Fig S5), and the proportion of DA proteins is similar (16% for DDA & 28% for DIA). These results suggest that protein abundance changes in response to DOX identified by DIA are replicated when abundance data is collected through data-dependent acquisition.

We identified a set of proteins with a high level of intra-modular connectivity that are likely to play a central role in the biological processes associated with each module. These 403 hub proteins have the highest correlation with the module eigenproteins and the highest intra-modular connectivity (kIN) score (top 10% of all network proteins; See Methods; Fig S6-7 & Table 2). Hub proteins predominantly reflect the module’s collective response to DOX or VEH treatment. For example, DDX27 in the α module is downregulated in response to DOX treatment, while PIGQ in the λ module shows no difference in abundance in response to DOX (Fig 2E). We then focused on the relationship between hub proteins across the five DOX-correlated modules (n = 202). The resulting network revealed not only strong intra-modular connections but also significant inter-modular correlations among proteins with a similar direction of effect in response to DOX treatment, indicating potential roles in inter-modular overlap for biological processes (Fig 2F).

### Module proteins differ in their tissue specificity and cellular localization

Having identified modules of co-expressed proteins that are correlated with DOX treatment, we next sought to investigate the properties associated with each module. To determine the specificity of module proteins to heart tissue, we utilized data from the Human Protein Atlas (HPA) and the Genotype-Tissue Expression (GTEx) projects (20, 33). These resources provide extensive gene and protein expression measurements across various tissues. The HPA database highlights 419 genes with elevated expression in heart tissue, defined as at least a four-fold higher mRNA level in the heart compared to the average in other tissues. We detect 277 proteins corresponding to heart-elevated genes in our data. Heart-elevated proteins constitute only a small proportion of proteins in each module regardless of DOX-correlation status (median all = 4.5%, range all = 0 - 8%; Fig 3A). For a quantitative analysis of tissue specificity of DOX-correlated module proteins, we obtained tissue specificity (TS) scores for proteins in heart left ventricle tissue from GTEx. DA proteins generally exhibit lower TS scores than non-DA proteins (Wilcoxon rank-sum test; *P* < 0.05; Fig S8), suggesting that iPSC-CM proteins, which are DA in response to DOX, show reduced specificity to heart ventricle tissue. We then asked whether this trend applies at the module level. For each module in the network, module TS scores were compared to the mutually exclusive set of all other network proteins (network TS Median = 0.33). We find significant deviations in TS scores from DOX-correlated modules α, β, γ, and δ, as well as non-DOX-correlated modules κ, and ι (Wilcoxon rank-sum test; *P* < 0.05). Modules β, δ, κ, and ι have higher TS scores, whereas α and γ have lower TS scores (Fig 3B). DOX-correlated modules (α, β, γ, δ, and ε) are therefore heterogeneous in their tissue specificity. However, the module with the strongest correlation to DOX, α, contains proteins that are the least specific to heart ventricle tissue (TS score = -0.465). These findings suggest that proteins that are most responsive to DNA damage in cardiomyocytes are not specific to the heart ventricle; however there are still likely to be tissue-specific effects among DOX response modules as indicated by the β and δ modules.

**Figure 3:**
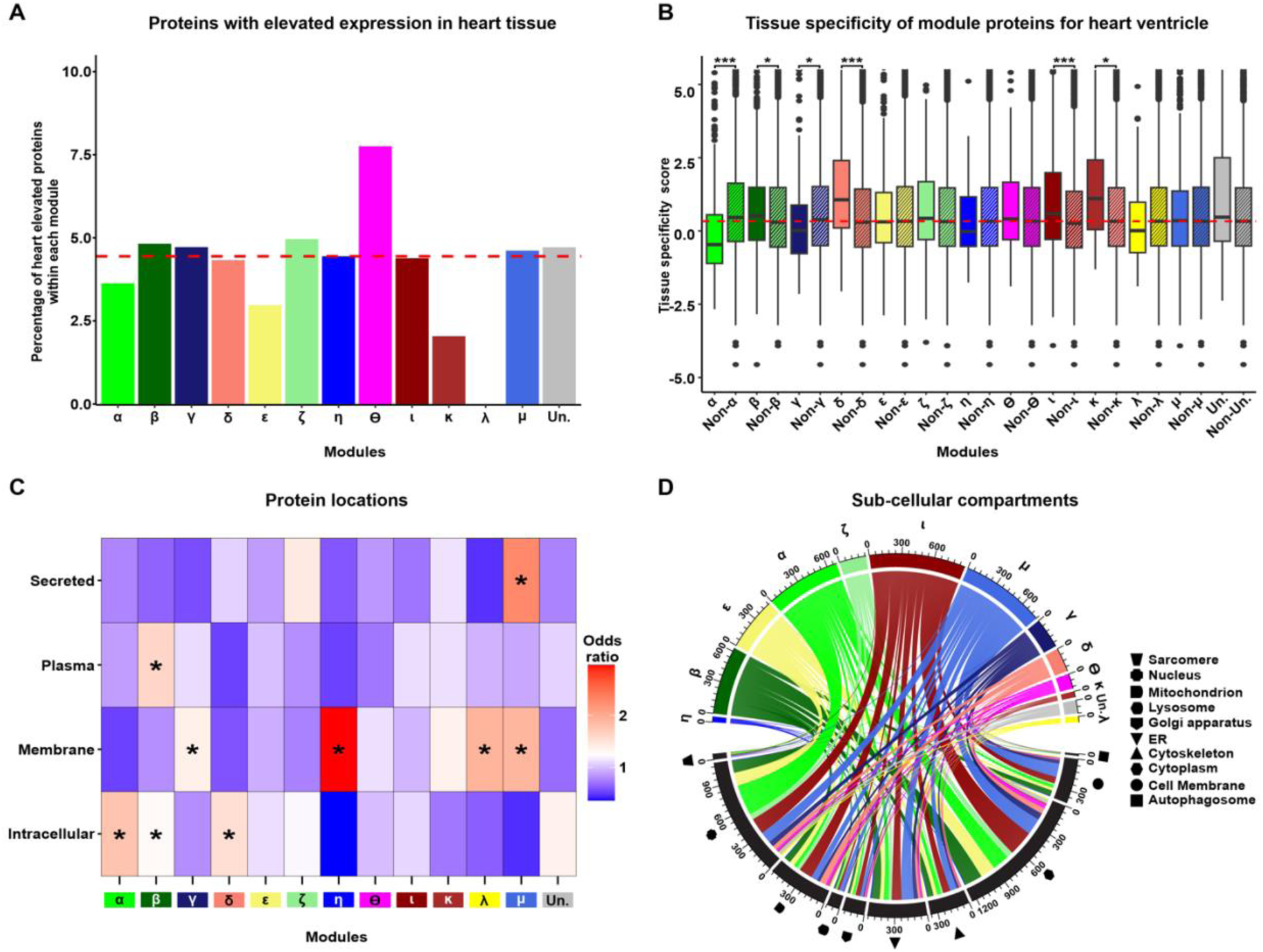
Network modules display heterogeneity for cellular localization and tissue specificity. **(A)** Percentage of each module in the network that consists of proteins annotated as having increased expression in heart relative to other tissues (20). Dashed red line represents the median percentage of heart elevated proteins across all modules. **(B)** Comparison of heart ventricle tissue specificity scores for each module (x) in the network compared to proteins in all other modules (Non-x). The red dashed line indicates the median tissue specificity score for the network. Asterisk represents statistically significant differences in tissue-specificity scores between module proteins and non-module proteins (\**P* < 0.01, \*\*\**P* < 0.0001). **(C)** Enrichment of module proteins across cellular and extracellular compartments. Asterisk represents locations with a significant enrichment of module proteins (\**P* < 0.05). **(D)** Distribution of module proteins across subcellular compartments. Colors represent the set of proteins in each module and shapes represent the cellular compartments. Numbers indicate the number of proteins in each module or compartment.

We next asked whether proteins within each module are restricted to specific intracellular or extracellular locations. We first investigated the localization of module proteins across four broad categories: intracellular, membrane-bound, plasma-detected, and secreted proteins as defined by HPA. We find that DOX-correlated modules α, β, and δ contain proteins that are enriched in the intracellular category compared to proteins not contained within each of these modules (Fig 3C; Fisher’s exact test; Odds ratio = 1.8, 1.3, 1.5 respectively; *P* < 0.05). Module γ is the only DOX-correlated module enriched for membrane proteins (Odds ratio = 1.4; *P* < 0.05), along with non-DOX-correlated modules η, λ and μ. Plasma-detected proteins are only enriched in module β (Odds ratio = 1.61; *P* < 0.05), while secreted proteins are only enriched in module μ (Odds ratio = 2.3; *P* < 0.05). The ε module is the only DOX-correlated module not enriched for any category. These results suggest that the cellular localization of proteins differs across modules. We then asked whether the sub-cellular localization of intracellular proteins differs across modules using annotation data from the UniProt database (34). Network proteins are generally distributed across multiple sub-cellular organelles including the sarcomere, nucleus, mitochondrion, lysosome, golgi apparatus, endoplasmic reticulum, cytoskeleton, cytoplasm, cell membrane, and autophagosome (Fig 3D). Nuclear proteins are enriched in modules α, β, and ε (*P* < 0.05). Modules α and ε also share enrichment for cell membrane, cytoplasm, endoplasmic reticulum, mitochondrial and lysosomal proteins. Module δ is enriched for mitochondrial proteins, while γ is enriched for endoplasmic reticulum and cell membrane proteins.

### DOX-correlated module proteins are enriched for distinct biological processes

To elucidate the broad functional roles of proteins within the DOX-correlated modules, we tested whether proteins associated with distinct biological processes are enriched in each module (See Methods). We found that the α, β, δ and ε modules show enrichment for various biological processes (Fisher’s exact test; adjusted *P* < 0.05; Fig 4A & Fig S9). The α module is enriched for 160 unique processes related to the DNA damage response, gene regulation via RNA splicing and metabolism, as well as chromatin regulating processes such as histone methylation and acetylation (Table 3). The β module is uniquely enriched for 35 processes related to protein localization within the nucleus and metabolism of carbohydrates, organophosphates, and oxoacids. Proteins in the δ module are uniquely enriched for 13 processes related to mitochondrial gene expression, translation and ATP synthesis. The ε module is uniquely enriched for 26 processes related to DNA replication and chromatin assembly. There are no processes significantly enriched within the γ module; however the most represented processes include glutamine family amino acid catabolic processes and ion homeostasis. Biological processes shared across modules include cytoplasmic translation (enriched in δ & ε), post-transcriptional regulation of gene expression (α & ε), and ribosome and ribonucleoprotein complex biosynthesis (α, ε & δ). Together, these data show a diversity of processes enriched amongst DOX-correlated modules.

**Figure 4:**
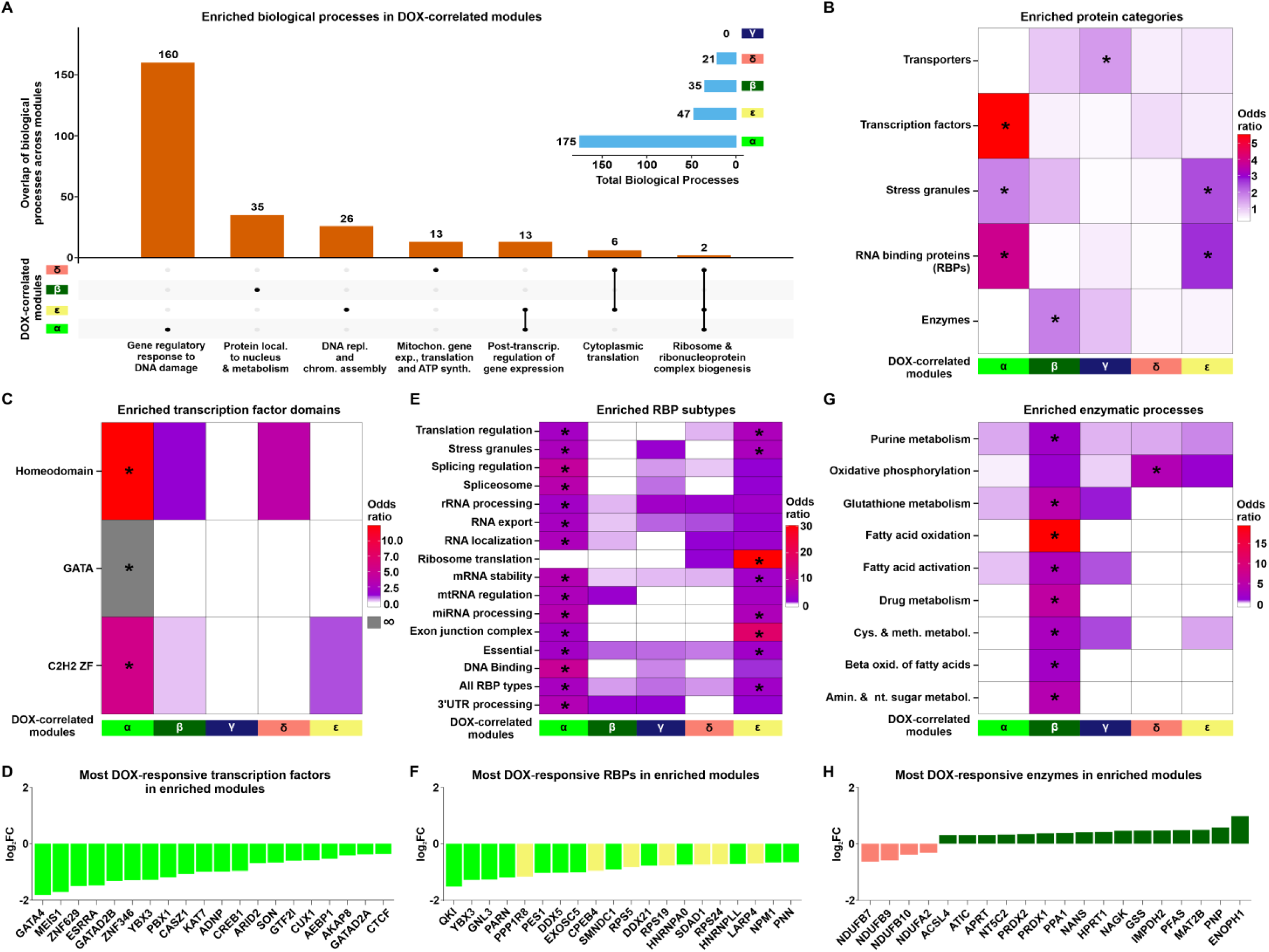
DOX-correlated module proteins are enriched for distinct functions. **(A)** The total number of enriched biological processes (adjusted *P* < 0.05) across DOX-correlated modules and their overlap between modules. Terms representing the top enriched processes are indicated. **(B)** Enrichment of proteins of different functional categories amongst DOX-correlated module proteins. Asterisk represents protein categories with a significant enrichment of module proteins (\**P* < 0.05). **(C)** Enrichment of proteins annotated by transcription factor binding domains amongst DOX-correlated module proteins. Grey shading indicates an infinite likelihood due to all transcription factor proteins with the particular binding domain being in only one module. **(D)** Top 20 most DOX-responsive transcription factors across DOX-correlated modules represented by log_2_ fold change from the pairwise differential abundance test. **(E)** Enrichment of RNA binding protein (RBP) types amongst DOX-correlated module proteins. **(F)** Top 20 most DOX-responsive RBPs across DOX-correlated modules. **(G)** Enrichment of proteins annotated by enzymatic process amongst DOX-correlated module proteins. **(H)** Top 20 most DOX-responsive enzymes across DOX-correlated modules.

We then asked whether proteins in each module are enriched for specific protein families consistent with their distinct biological processes. The α module is most strongly enriched for the splicing factor SR family, RNase PH family, and Histone deacetylase family (Fisher’s exact test; *P* < 0.05; Fig S10). β is most enriched for the TCP-1 chaperonin family, Periredoxin family, and Eukaryotic mitochondrial porin family. γ is enriched for proteins in the Septin GTPase family, AGC Ser/Thr protein kinase family, and Peptidase C2 family. δ is enriched for proteins in the 14-3-3 family, Adaptor complexes small subunit family, and Troponin I family. ε is enriched for proteins in the fibrillar collagen family, Ruvb family, and Lin-7 family. These results generally corroborate the biological process enrichment analysis results, while also underscoring that although some modules may overlap in terms of their biological processes, the proteins that are engaged in those processes across different modules can be differentiated by their protein families.

We selected five key protein categories to investigate further based on our module-specific functional enrichment results (Fig 4B; See Methods) (20, 35–37). The α module is uniquely enriched for transcription factors (Fisher’s exact test, OR = 5.6; *P* < 0.05; Fig 4C), particularly the homeodomain, GATA and C2H2 Zinc Finger families that include the GATA4, MEIS1 and ZNF629 transcription factors (Fig 4D). Mutations in GATA4 are associated with atrial septal defects, arrhythmia, and a reduced capacity for the cardiac hypertrophic response (38), and have been implicated in DOX-induced cardiotoxicity (39). Both the α and ε modules show enrichment for stress granule components and RNA Binding Proteins (*P* < 0.05; Fig 4B). RNA binding proteins related to multiple post-transcriptional processes including splicing, translation and mRNA stability are enriched (Fig 4E), and include proteins such as QKI, YBX3 and GNL3 (Fig 4F). QKI regulates pre-mRNA splicing, export of mRNAs from the nucleus, protein translation, and mRNA stability (40), and is implicated in cardiomyocyte calcium dynamics and contractility (41), cardiomyopathies (40) and attenuation of DOX-induced cardiotoxicity (42). The β module uniquely exhibits enrichment for enzymes (OR = 1.9; *P* < 0.05; Fig 4B) involved in processes such as fatty acid oxidation, glutathione metabolism and purine metabolism (Fig 4G). The most DOX-responsive enzymes include NDUFB7, a member of the δ module and critical component of the mitochondrial membrane respiratory chain complex I, and ENOPH1 a member of the β module, and key enzyme in the methionine salvage pathway (Fig 4H). The γ module is distinctively enriched for transporter proteins including PRDX2 and ABCB1 (OR = 1.6; *P* < 0.05; Fig 4B). These results are consistent with the biological processes enriched in each module.

### Hub proteins in DOX-correlated modules are depleted for pQTLs

We next focused our attention on the properties of the 403 hub proteins within the network. We were particularly interested in understanding the tolerance of these central proteins to physiological genetic variation. To do so, we investigated proteins whose expression level varies across individuals depending on the genotype of an associated SNP i.e. protein quantitative trait loci (pQTLs; Fig 5A). We obtained a set of pQTLs identified in human plasma from thousands of individuals (17), and overlapped these with iPSC-CM network proteins (Table 2). We first asked whether pQTL proteins associated with either *cis-* or *trans*-SNPs are enriched amongst hub proteins. Hub proteins are nether enriched nor depleted for pQTLs (Fisher’s exact test OR = 1.1; 95% CI = 0.8-1.5 for *cis*-pQTLs and OR = 1.0; 95% CI = 0.8-1.4 for *trans*-pQTLs). We then asked whether pQTL proteins are enriched amongst proteins that respond to DNA damage. DOX-correlated proteins are modestly depleted for pQTL proteins associated with either *cis-* (OR = 0.7; 95% CI = 0.6-0.8; *P* < 0.05; Fig 5B) or *trans*-SNPs (OR = 0.8; 95% CI = 0.7-1.0; *P* < 0.05, Fig 5B). However, DOX-correlated hub proteins show an even more pronounced depletion for pQTL proteins mapped to *cis-* (OR = 0.2; 95% CI = 0.1-0.5; *P* < 0.05; Fig 5B) or *trans*-SNPs (OR = 0.4; 95% CI = 0.2-0.6; *P* < 0.05; Fig 5B).

**Figure 5:**
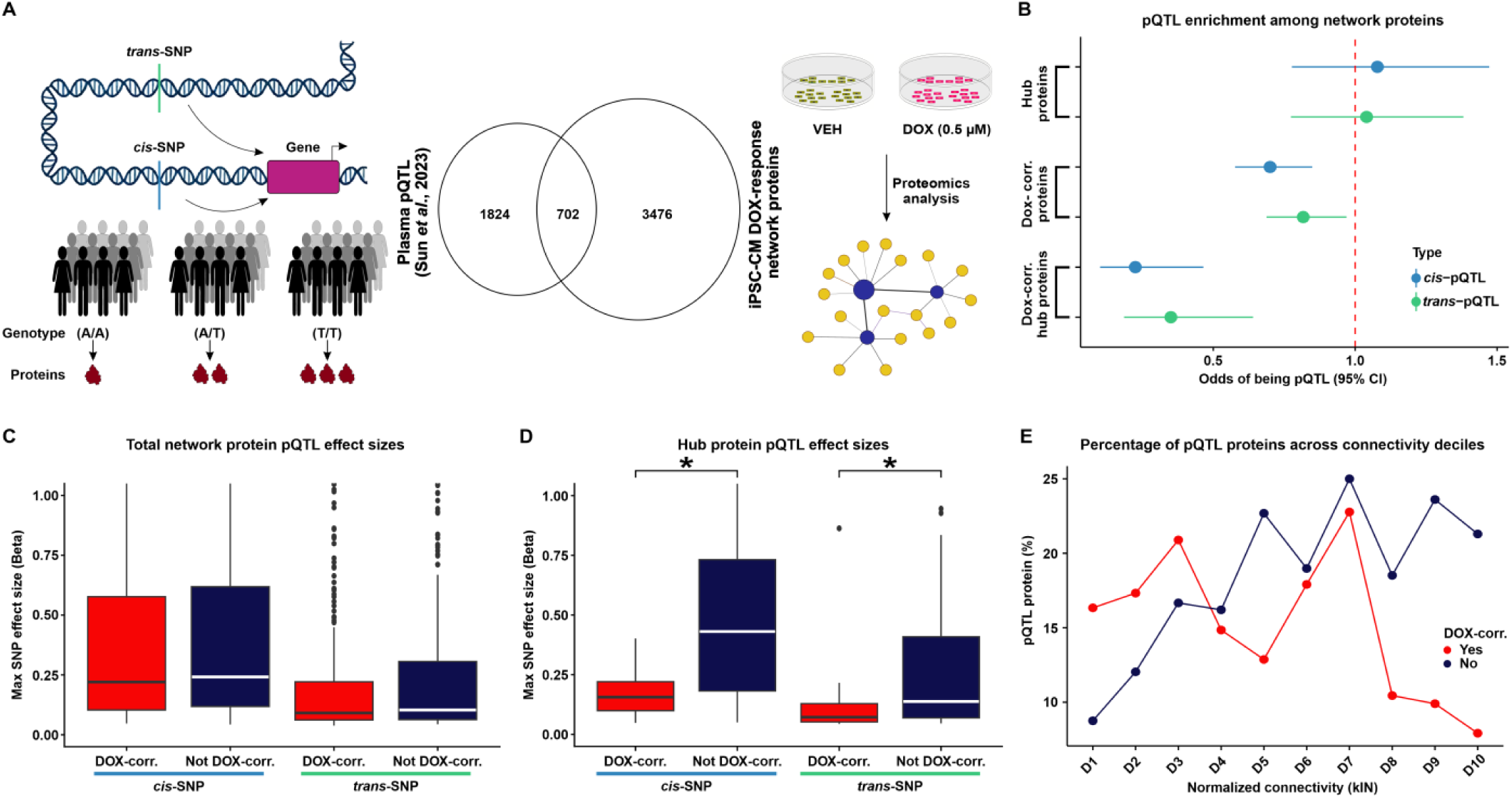
DOX-correlated hub proteins are depleted for protein quantitative trait loci. **(A)** Schematic representing the rationale behind the test design. Protein expression can be influenced by *cis*- or *trans*-SNPs resulting in differential abundance of proteins across individuals with different genotypes. We integrated existing protein quantitative trait loci (pQTL) data from plasma samples (17) with our DOX-treated iPSC-CM protein network. **(B)** Enrichment of *cis*- (blue) and *trans*-pQTLs (green) amongst hub proteins, DOX-correlated proteins, and DOX-correlated hub proteins. **(C)** Maximum pQTL effect sizes for *cis*- and *trans*-pQTLs amongst all proteins in DOX-correlated modules and non-DOX-correlated modules. **(D)** Maximum pQTL effect sizes for *cis*- and *trans*-pQTLs amongst all hub proteins in DOX-correlated modules and non-DOX correlated modules. Asterisk represents a significant difference in effect sizes between DOX-correlated and non-DOX-correlated modules (\**P* < 0.05). **(E)** Percentage of pQTL proteins across connectivity deciles for normalized kIN for proteins in DOX-correlated modules (red), and non-DOX-correlated (blue). Deciles are ordered by increasing kIN scores.

Next, we investigated those pQTLs that correspond to proteins expressed within our network and asked whether the SNP effect size of the pQTLs differed across network components. We assigned each pQTL protein to the *cis-* or *trans*-SNP with the greatest effect size. As expected, the median effect size for *cis*-pQTL proteins in the network is greater than that for *trans*-pQTLs (0.2 vs. 0.1). There is no difference in the *cis*- or *trans*-pQTL effect sizes between DOX-correlated and non-DOX-correlated proteins (Fig 5C). However, DOX-correlated hub proteins have lower *cis-* and *trans*-pQTL effect sizes than hub proteins that are not correlated with DOX (Wilcoxon rank-sum test; *P* < 0.05; Fig 5D).

Given the depletion of pQTLs amongst DOX-correlated hub proteins and not the total set of hub proteins, we next asked if there is a relationship between the intra-modular connectivity of a protein and the probability of that protein being a pQTL that is influenced by DOX. We assigned proteins into two groups based on their DOX-correlation status, stratified proteins within each group into deciles based on their connectivity, and calculated the percentage of pQTL proteins in each decile. In the non-DOX-correlated group, the decile with the highest connectivity shows the greatest percentage of pQTL proteins (21%), while the decile with the lowest connectivity shows the lowest percentage of pQTL proteins (9%). Across deciles, there is a general upward trend in the percentage of pQTL proteins as intra-modular connectivity increases (Fig 5E). In the DOX-correlated group, the highest connectivity decile shows a reduction in the percentage of pQTL proteins (8%) relative to the lowest connectivity decile (16%), and more variability for the percentage of pQTL proteins across deciles. Therefore, the percentage of pQTL proteins across different deciles showed opposite trends depending on their correlation to DOX. The decrease in pQTL proteins with the highest connectivity among DOX-correlated proteins suggests that hub proteins correlated to DOX treatment are under stronger evolutionary constraints, leading to reduced genetic variation in these highly connected proteins. In summary, our analysis reveals that DOX-correlated hub proteins are less likely to be associated with pQTLs, indicating their likelihood to play an essential role in maintaining network stability and function under stress conditions.

### DOX-correlated hub proteins are enriched for loss-of-function intolerant proteins

Given that DOX-correlated hub proteins are depleted for proteins that vary in their expression across healthy individuals, we next asked about the tolerance of these proteins to variation more broadly. We utilized several genetic tolerance scores to assess the functional impact and evolutionary constraint of genes encoding proteins within specific modules. First, we considered the probability of each protein in a module being haploinsufficient (pHaplo), which estimates whether a single copy of a gene can maintain normal function, or triplosensitive (pTriplo), which estimates the risks of gene dose increases, that can be equally detrimental. Genes with a high pHaplo score (≥ 0.86) or a high pTriplo score (≥ 0.94) are deemed haploinsufficient or triplosensitive, respectively, and likely precipitate health consequences (43). We compared the gene dose scores of DOX-correlated module proteins against the broader network (See Methods). The α and ε modules have higher pHaplo scores compared to network proteins, while δ module proteins have lower pHaplo scores (Fig 6A; Wilcoxon rank-sum test; *P* < 0.05). The α module also shows increased pTriplo scores compared to the network average, while the δ module’s pTriplo scores are lower (Fig 6B; *P* < 0.05). These results show that the most DOX-correlated module, α, is most sensitive to gene dosage changes.

**Figure 6:**
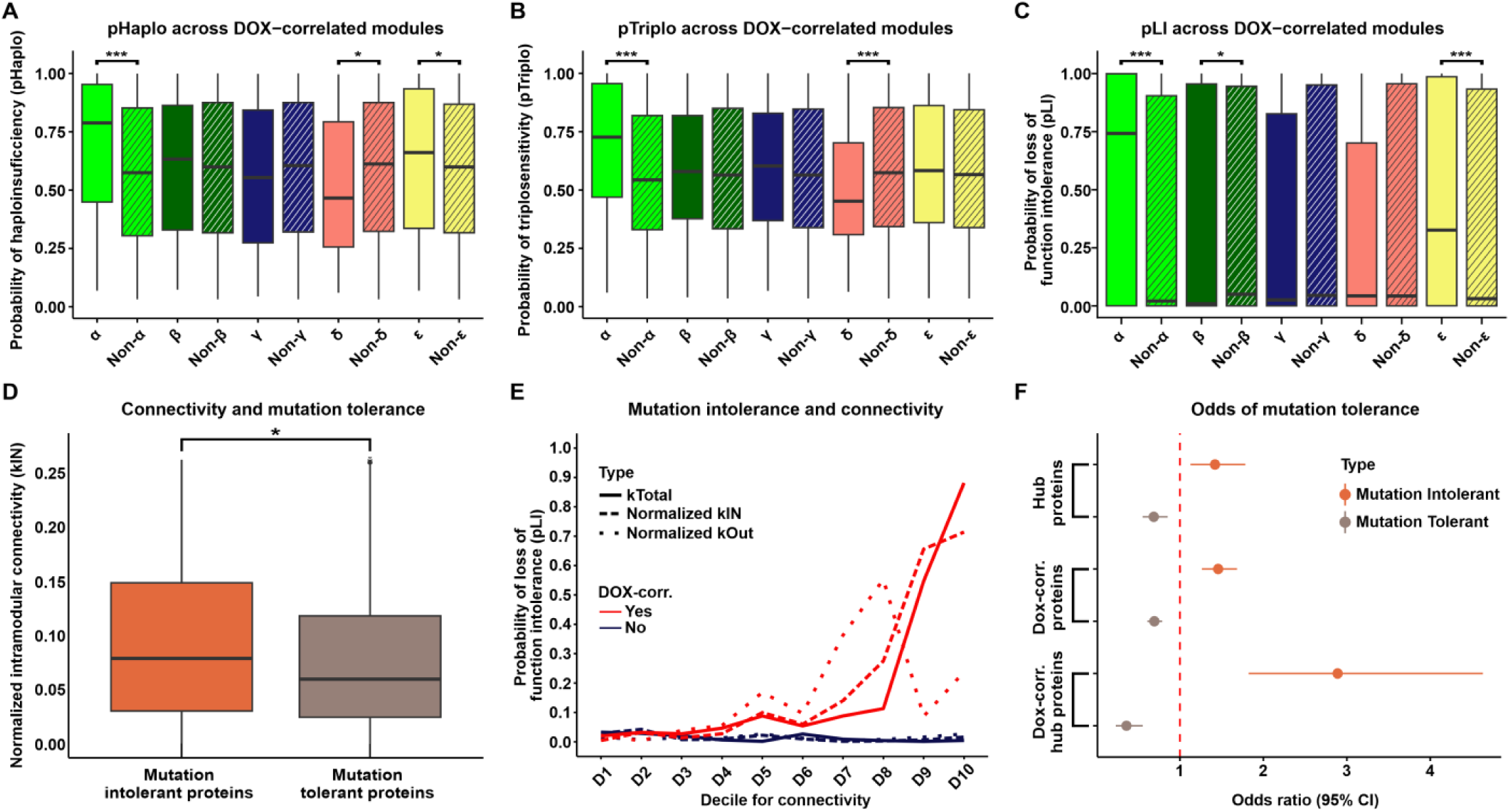
DOX-correlated hub proteins are enriched for loss-of-function intolerant proteins. **(A)** Probability of haploinsufficiency (pHaplo) for all proteins in each DOX-correlated module (x), and all proteins outside of the module (Non-x). Asterisk represents a significant difference in scores between module-specific proteins and all proteins outside of modules (\**P* < 0.01, \*\*\**P <* 0.0001). **(B)** Probability of triplosensitivity (pTriplo) for all proteins in each DOX-correlated module (x), and all proteins outside of the module (Non-x). **(C)** Probability of loss-of-function intolerance (pLI) for all proteins in each DOX-correlated module (x), and all proteins outside of the module (Non-x). **(D)** Connectivity (normalized kIN) amongst mutation tolerant (pLI ≤ 0.1) and mutation intolerant (pLI ≥ 0.9) proteins. Asterisk represents a significant difference in scores between module-specific proteins and all proteins outside of modules (\**P* < 0.05). **(E)** Median pLI scores across proteins in connectivity deciles (kTotal (solid line), normalized kIN (dashed line), normalized kOut (dotted line)) for proteins in DOX-correlated modules (red), and non-DOX-correlated (blue). Deciles are ordered by increasing connectivity scores. **(F)** Enrichment of mutation intolerant (orange) and mutation tolerant (brown) proteins for hub proteins, DOX-correlated proteins, and DOX-correlated hub proteins.

Next, we examined the tolerance of module proteins to mutations that reduce or eliminate protein function. Genes with a high probability (≥ 0.9) of loss-of-function intolerance (pLI) are considered loss-of-function intolerant, meaning they cannot withstand loss-of-function mutations without significant phenotypic impact. Conversely, genes with a low probability (≤ 0.1) can tolerate loss-of-function mutations with minimal phenotypic consequences and are considered loss-of-function tolerant (43, 44). Both α and ε module proteins exhibit significantly higher pLI scores than proteins outside these modules (Fig 6C; Table 2; *P* < 0.05), indicating that they are less tolerant to genetic variation. In contrast, the β module is characterized by lower pLI scores (Fig 6C; *P* < 0.05), suggesting these proteins can better tolerate loss-of-function mutations. These results emphasize that the α module contains proteins that are highly intolerant to pathogenic variation.

To test the relationship between modular connectivity and pLI across the network, we identified the set of loss-of-function tolerant and loss-of-function intolerant proteins. We find that loss-of-function intolerant proteins have a higher level of intra-modular connectivity compared to loss-of-function tolerant proteins (Wilcoxon rank-sum test; *P* < 0.05; Fig 6D). To understand the relationship between different types of connectivity and DOX-correlation, we next considered the spectrum of tolerance to mutation across all proteins in the network. First, we classified all proteins into two groups based on their DOX-correlation status. We then generated deciles for intra-modular connectivity (kIN), inter-modular connectivity (kOut) and total network connectivity (kTotal) for both DOX-correlated, and non-DOX-correlated proteins. For DOX-correlated proteins, there is a marked increase in pLI scores in the highest kTotal and kIN deciles (Fig 6E). Conversely, pLI scores for kOut appeared more variable at higher connectivity deciles. In comparison to DOX-correlated proteins, median pLI scores for non-DOX-correlated proteins remain consistently low across deciles for all measures of connectivity. These findings highlight that DOX-correlated proteins with the highest total connectivity, primarily driven by intra-modular connectivity, are essential for cellular function under DNA damage conditions and are under strong evolutionary constraints to maintain their functional integrity. In contrast, non-DOX-correlated proteins do not exhibit a relationship between connectivity and mutation intolerance, suggesting a less critical role in maintaining network stability.

We next asked whether loss-of-function intolerant proteins are enriched among proteins central to the protein network given that there are increased pLI scores in the highest decile for intra-modular connectivity. DOX-correlated proteins are modestly enriched for proteins whose encoding genes are loss-of-function intolerant (Fisher’s exact test; OR = 1.5; CI = 1.3-1.7; *P* < 0.05; Fig 6F) and depleted for proteins that are tolerant to mutation (OR = 0.7; CI = 0.6-0.8; *P* < 0.05). Hub proteins are modestly enriched for loss-of-function intolerant proteins (OR = 1.4; CI = 1.1-1.8; *P* < 0.05) and depleted for mutation tolerant proteins (OR = 0.7; CI = 0.6-0.9; *P* < 0.05). However, DOX-correlated hub proteins are highly enriched for mutation-intolerant proteins (OR = 2.9; CI = 1.8-4.6; *P* < 0.05) and depleted for mutation-tolerant proteins (OR = 0.4; CI = 0.2-0.6; *P* < 0.05; Fig 6F). These data demonstrate that DOX-correlated hub proteins are likely to be the most critical proteins to the DNA damage response network.

### DOX-correlated modules are enriched for cardiovascular disease-associated proteins

We assessed the relative likelihood for DOX-correlated proteins to be contained in the set of proteins mapped to cardiovascular traits. Because our network’s modules are enriched for biological processes and protein families, testing for enrichment of proteins in trait-associated loci within network modules can contextualize how genetic risk factors might disrupt specific biological pathways and contribute to disease. We therefore collated heart function measurement traits and CVD traits from the GWAS catalog (45), and tested for enrichment of the GWAS mapped genes amongst proteins expressed in our network modules (Fig 7A; See Methods). We obtained 1,077 traits to explore, where 993 correspond to heart function measurements, and 84 correspond to CVDs. DOX-correlated modules are enriched for proteins mapped to 26 clinical cardiovascular traits (Fisher’s exact test; *P* < 0.05), 25 of which are unique to DOX-correlated modules, and five CVDs, all of which are specific to a module (Fig 7B).

**Figure 7:**
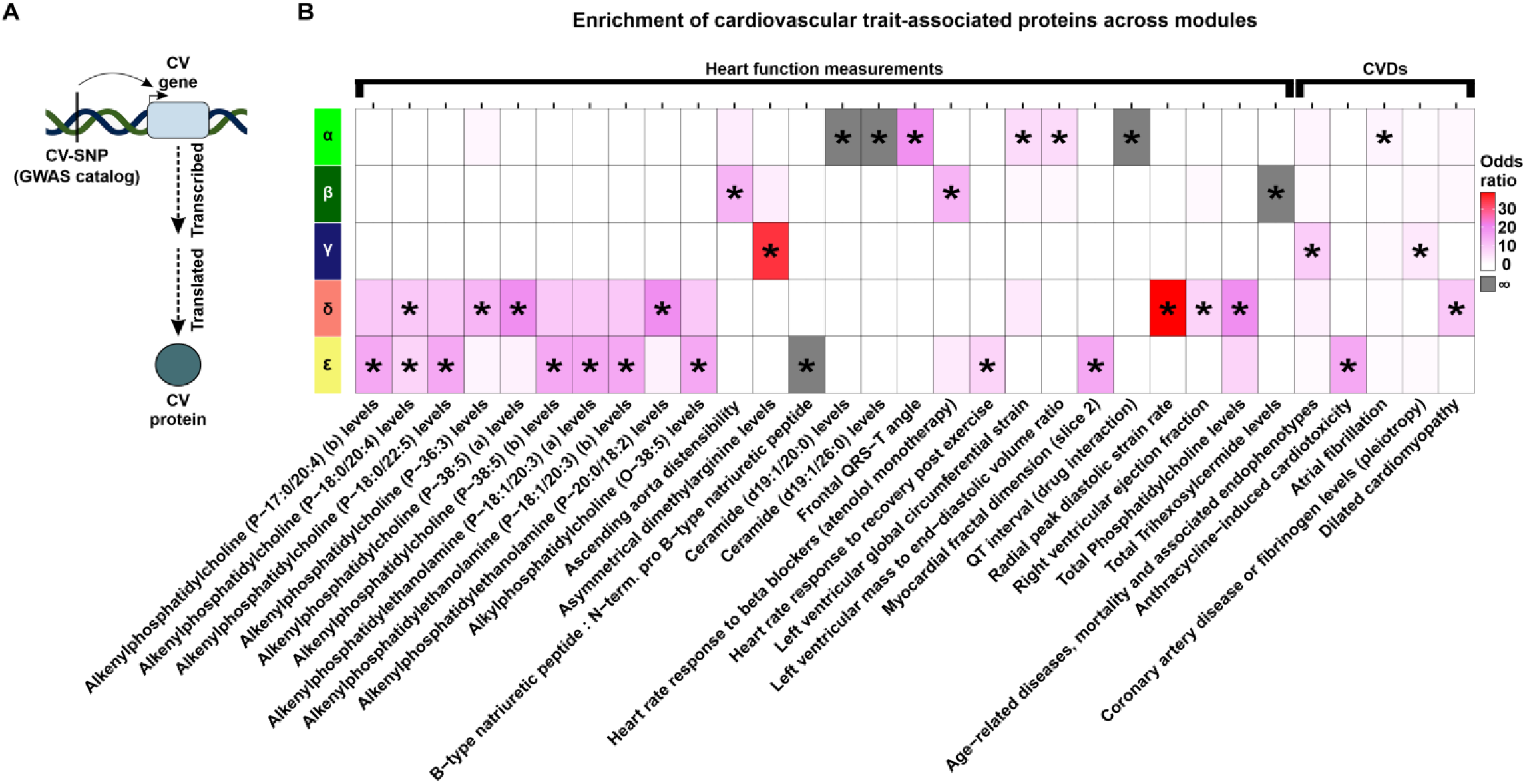
DOX-correlated modules are enriched for proteins mapped to cardiovascular traits and diseases. **(A)** Schematic representing how cardiovascular trait proteins are obtained. Cardiovascular trait-associated SNPs (CV-SNP) from GWAS are mapped to nearby genes (CV gene) and converted to the corresponding protein identifier (CV protein). **(B)** Enrichment of heart function measurement traits (GWAS catalog ontology class EFO_0004311) and heart diseases (EFO_0003777) across DOX-correlated module proteins. Grey shading indicates an infinite likelihood of enrichment due to all trait-associated proteins being in only one module. Asterisk represents traits and diseases that are enriched amongst DOX-correlated module proteins (\**P* < 0.05) and contain at least two proteins in the module.

The α module is enriched for proteins related to clinical traits such as QT-interval, left ventricular mass to end diastolic volume ratio, frontal QRS-T angle, and ceramide levels. These proteins are also relevant to disease as this module contains proteins implicated in atrial fibrillation. The β module is enriched for trihexosylceramide levels, heart rate response to beta blockers, and ascending aorta distensibility. However, the β module is not enriched for any CVD traits. The γ module is enriched for age-related endophenotypes (classified as both a clinical trait and CVD) and asymmetrical dimethylarginine levels. The δ module is enriched for glycerophospholipid-related traits as well as radial peak diastolic strain and right ventricular ejection fraction. Proteins in this module are enriched for dilated cardiomyopathy. The ε module is enriched for traits related to glycerophospholipids, myocardial dimensions, heart rate post exercise, and the ratio of BNP to pro-BNP. This module is also enriched for proteins associated with Anthracycline-induced cardiotoxicity. Together these results highlight that the modules of co-expressed proteins that we identify are associated with distinct physiological and pathological processes, suggesting protein co-expression may serve as a functionally relevant context to interpret the role of CVD risk proteins in mediating CVD.

### DOX-correlated hub proteins are enriched for physical protein-protein interactors of CVD-associated proteins

We assessed the relative likelihood for key network proteins to be contained in the set of proteins mapped to CVD (Fig 8A). CVD-associated proteins are neither enriched nor depleted amongst hub proteins, DOX-correlated proteins or DOX-correlated hub proteins (Fig 8B) suggesting that they do not contribute to the key features of our protein network. Although these results are in line with the observation that DOX-correlated hub proteins are depleted for pQTLs and are intolerant to genetic mutation, these data conflict with the result showing enrichment of proteins associated with CVD within DOX-correlated modules.

**Figure 8:**
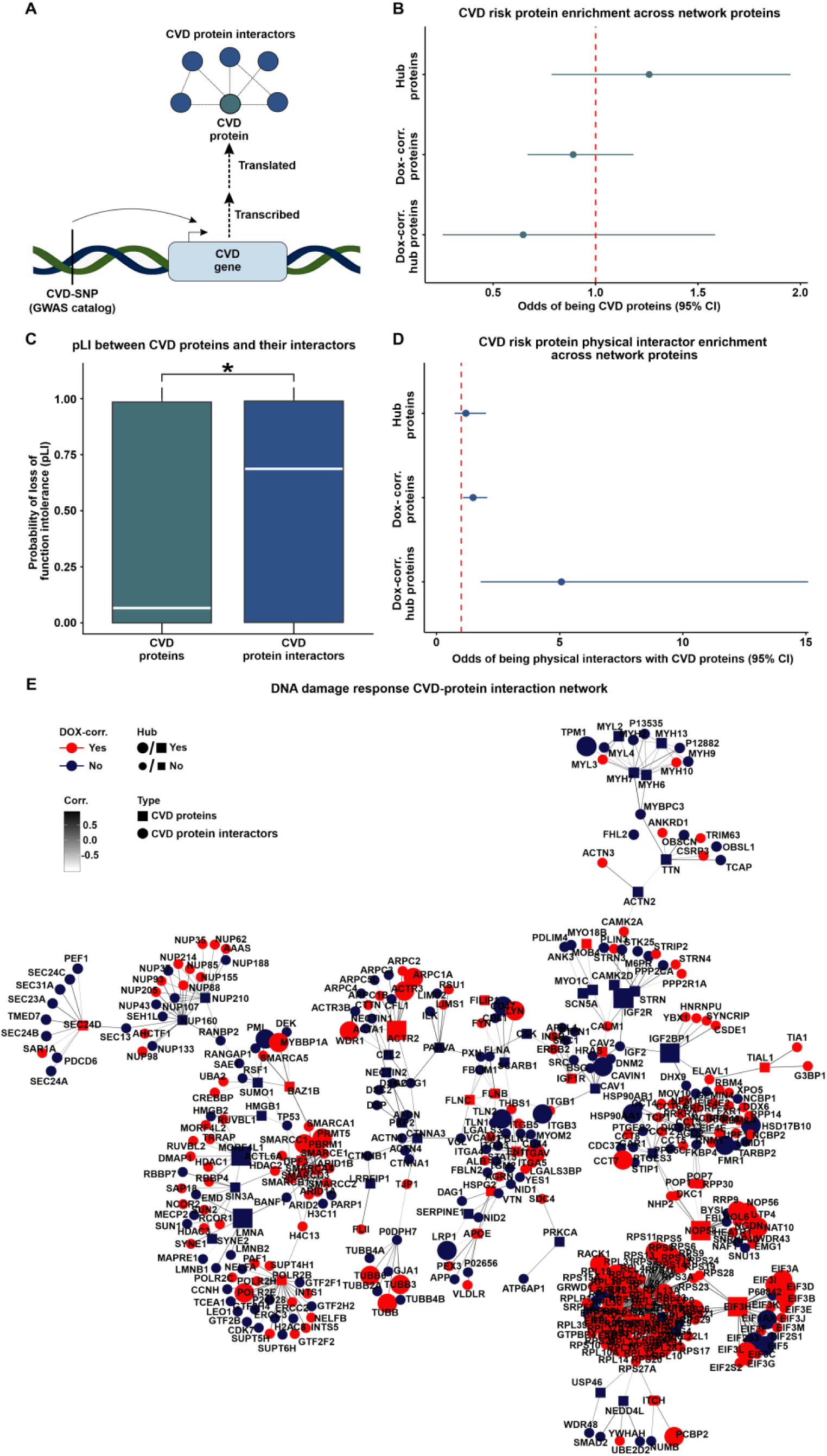
DOX-correlated hub proteins are enriched for physical protein interactors of CVD risk proteins. **(A)** Schematic representing the rationale behind the test design. CVD-associated SNPs (CVD-SNP) are mapped to nearby genes (CVD gene) which are translated into proteins (CVD proteins) that may physically interact with other proteins (CVD protein interactors). **(B)** Enrichment of CVD risk proteins amongst hub proteins, DOX-correlated proteins, and DOX-correlated hub proteins. **(C)** pLI score distribution of CVD risk proteins and CVD risk protein interactors. Asterisk represents a significant difference in the scores between protein groups (\**P* < 0.05). **(D)** Enrichment of CVD risk protein interactors amongst hub proteins, DOX-correlated proteins, and DOX-correlated hub proteins. **(E)** Protein-protein interaction network for CVD proteins (square) and CVD protein interactors (circle) expressed within the co-expression network. Color represents if the protein is in a DOX-correlated module (red) or a non-DOX-correlated module (blue). Edges represent weighted correlation within the co-expression network. Node size indicates if a protein is a hub (large icon) or not a hub (small icon) protein. A CVD protein subnetwork containing the most highly connected proteins is presented here with the full CVD protein network displayed in Fig S12.

We next reasoned that proteins that are known to physically interact with CVD-associated proteins may provide insight into how proteins encoded by CVD risk genes connect to the DOX-correlated proteins in our network. We therefore identified pairwise interactions between all 4,178 network proteins (STRINGdb; confidence score ≥ 0.9) and generated a total physical protein-protein interaction (PPI) network (n = 13,750 edges). We then extracted the subnetwork containing CVD proteins (n = 213) and their interactors (n = 798; n = 909 edges; Tables 2 & 4). We then asked whether CVD proteins and their interactors differ in their essentiality. CVD proteins have lower pLI scores than proteins they interact with (0.07 vs. 0.69; Wilcoxon rank-sum test; *P* < 0.05; Fig 8C). To test the robustness of these results, we analyzed 10,000 randomly generated subnetworks from the total network that maintained a similar degree distribution as the CVD protein network (See Methods). The enrichment *P* value of CVD-associated proteins and their interactors is lower than the 5^th^ percentile of the random distribution (*P* = 0.00001 vs. *P* = 0.006), and the median pLI score difference between our test sets (0.62) is within the 95^th^ (0.63) and 96^th^ (0.64) percentiles of the randomly generated networks. These findings suggest that genes encoding CVD risk proteins are less likely to be essential compared to their interacting partners.

We next asked about the relative likelihood for hub proteins in the network to be contained in the set of CVD protein interactors. We first calculated the frequency of protein interactors falling in the same module. We find that the proportion of interacting proteins expressed in the same module is higher for interactions where one of the proteins is a hub protein compared to interactions that do not include a hub protein (Wilcoxon rank-sum test; *P* < 0.05; Fig S11). Although hub proteins are more likely to be co-expressed in the same module with their physical protein interactors, hub proteins are not enriched among CVD protein interactors (Fig 8D). However, CVD protein interactors are enriched for DOX-correlated proteins in comparison to proteins that are not DOX-correlated (Fisher’s exact test, Odds ratio = 1.5, CI = 1.1-2.1; *P* < 0.05; Fig 8D). We further find that CVD protein interactors are enriched for DOX-correlated hub proteins in comparison to hub proteins that are not DOX-correlated (Odds ratio = 5.1; CI = 1.8-15.1; *P* < 0.05; Fig 8D). We similarly observe a significant enrichment of CVD protein interactors amongst DOX-correlated hub proteins when identifying interactors with a lower stringency score (n = 49,020 interactions; STRINGdb confidence score ≥ 0.4; Odds ratio = 2.3, *P* < 0.05). These data suggest that the pathogenicity of CVD variants identified by GWAS might not only be a consequence of a direct effect on a protein’s function but also indirectly through interactions with essential proteins that are highly correlated to the DNA damage response in cardiomyocytes. We therefore illustrate the CVD protein and CVD protein interactor network together with our experimentally-derived DOX-correlation and hub protein status annotations (Fig 8E & Fig S12).

We posit that the DNA damage-associated CVD protein network in cardiomyocytes can also be used to reduce the search space for druggable targets that might have the greatest impact on influencing various CVD phenotypes. We therefore also annotated each network protein by whether they are a target of an FDA-approved drug (Table 2) (33). We identified 79 proteins in DOX-correlated modules that are druggable including the chromatin-modifying enzymes HDAC1, HDAC2 and HDAC3. Seven DOX-correlated proteins are CVD proteins (FADS1, FADS2, FINC, IGF1R, AT2B1, RL3, and TRXR1), and five have elevated expression in heart tissue (RYR2, ADT1, LDHB, ODO1 and AAPK2), thereby opening further avenues of investigation.

## Discussion

Many genetic loci have been associated with CVD. While the genes in these loci can be inferred to play a role in disease risk, the mechanisms behind these associations is often unclear. We hypothesized that a relevant environmental factor may provide insight. DNA damage is a ubiquitous stressor that is both implicated in, and predictive of CVD (3). It can be induced through exogenous factors including through administration of drugs used in the treatment of cancer. To understand how DNA damage in cardiomyocytes influences CVD risk, we used an *in vitro* model of iPSC-CMs from multiple individuals to study the effects of DOX on the proteome. We identified many changes in protein abundance that can be connected to CVD risk proteins.

### DOX induces changes to the cardiomyocyte proteome relevant to anthracycline-induced cardiotoxicity

The effects of genetic variation and environmental perturbations on mediating risk to complex disease have most often been assayed through the transcriptome. This includes studies investigating the influence of DOX on cardiomyocytes (24, 27, 46). Here, we measured the effects of DOX on the cardiomyocyte proteome using both a pairwise differential abundance test, and a co-expression network correlated with DOX treatment. We identified 1,167 differentially abundant proteins (28% of the proteome), and five out of 12 co-expressed modules to be correlated with DOX treatment. A previous study that assayed the proteomic response to anthracyclines, including DOX, using both human microtissues, and tissues from heart failure patients identified four anthracycline-associated hub proteins (47). We find two of these proteins, BAG3 and CAND1, in DOX-correlated modules in cardiomyocytes indicating the relevance of our cardiomyocyte model. In addition, our study generated protein data for thousands more proteins implicating many more proteins in DOX toxicity. Conversely, we found little overlap between our data and DOX-treated rodent cardiac proteomic data from rat cardiomyocytes (48) and rat heart tissue (49) suggesting species-specificity in DOX responses.

Our network analysis identified the α module as the most DOX-correlated co-expression module. The 579 proteins in this module decrease in their abundance following DOX treatment and are enriched for proteins in gene regulatory processes and DNA damage repair. The most enriched processes relate to RNA processing and splicing. The associated proteins likely contribute to the large-scale splicing changes that have previously been identified following DOX treatment (27). Processes related to chromatin organization and histone modifications are also enriched, in line with previous work in a murine DOX toxicity model indicating effects on histone eviction and histone modifying enzymes (50). This module is also enriched for transcription factors including GATA4, CTCF, ZNF629, ESSRA, GATAD2B, ZNF346, YBX3, GTF2I, GATAD2A, indicating potential drivers of the gene expression changes observed due to DOX treatment in heart cells (38, 39, 51–61). Many of these processes are important across cell types, and we correspondingly observe decreased heart tissue specificity for proteins in this module. Indeed, DOX can lead to neurotoxicity, hepatotoxicity, and nephrotoxicity in addition to cardiotoxicity (62–64). Conversely the DOX-correlated δ module includes proteins with higher heart specificity than proteins in other modules, and is enriched for processes related to mitochondrial functions such as oxidative phosphorylation. Proteins in this module may thus contribute to some of the *in vivo* tissue-specific effects of DOX on the heart.

The target of DOX is TOP2, expressed in the heart as both the α and β isoforms. It is TOP2B that is thought to mediate the cardiotoxic effects of anthracycline chemotherapeutics such as DOX. While we do not detect a change in TOP2B protein abundance in response to DOX, it is co-expressed in the ε module, which is DOX-correlated and consists of proteins downregulated in response to DOX. Notably, ∼75% of proteins shown to physically interact with TOP2B (10/13 expressed in iPSC-CMs), including CTCF, are present in DOX-correlated modules α and ε and exhibit high connectivity (65). GWAS have identified 10 risk loci associated with anthracycline-induced cardiotoxicity (66–68). We find three proteins encoded by genes mapped to these risk loci expressed in our network, where two are enriched in the ε module. These proteins include POLRMT and RPL7. Decreased abundance of POLRMT, a DNA-directed RNA polymerase located in mitochondria, and RPL7, a component of the large ribosomal subunit, suggest that these proteins concordantly lead to decreased mitochondrial transcription and translation due to DOX treatment. While the mitochondrial topoisomerase TOP1MT is not detected in our network, TOP2A and TOP2B can translocate to mitochondria and resolve torsional stress of mitochondrial DNA (69–71). It stands to reason that POLRMT, a physical protein interactor of TOP1MT, may interact with TOP2B in the mitochondria under DNA-damaging conditions. Together, the proteins that we identify as responding to DOX in cardiomyocytes have been implicated in DOX-induced cardiotoxicity through molecular and genetic approaches.

### DOX-induced protein expression changes are relevant to CVD

Our network approach allowed us to intersect sets of DOX-correlated proteins with proteins implicated in risk for all complex CVDs. The α module is uniquely enriched for proteins encoded by genes in loci associated with atrial fibrillation. These proteins include GATA4, LRRC10, RBM20, SLIT3, CAND2, GTF2I, HSPG2, FILIP1, MYO18B, CASZ1, and DPF3 that are downregulated in response to DOX treatment. Many of these proteins show evidence for independently impacting atrial fibrillation risk, as well as playing an essential role in cardiac development (38, 39, 51–61). The co-expression of these proteins involved in the genetic risk for atrial fibrillation indicates that DNA damage may mediate atrial fibrillation by collectively reducing the abundance of these critical proteins. Atrial fibrillation can be a cause or consequence of many complex CVD traits (72), highlighting the importance of these proteins to disease.

DOX-correlated module δ captures proteins intersecting DNA damage and mitochondrial function, as well as risk proteins for dilated cardiomyopathy. For example, FHOD3 has been shown to play a critical role in regulating myocardial thickness in both dilated cardiomyopathy and hypertrophic cardiomyopathy progression (72–74). BAG3 haploinsufficiency disrupts many cardiomyocyte processes such as sarcomere integrity, apoptosis, calcium homeostasis, as well as mitophagy by interactions with E3 ubiquitin-protein ligase, which is co-expressed with BAG3 within the δ module (75).

In addition to providing insight into how specific proteins contribute to CVD risk, our study also has the potential to inform on proteins that might mediate cardioprotective effects to DOX. Metabolism-mediating therapies such as SGLT2 inhibitors are being clinically investigated for their role in cardioprotection following heart failure and anthracycline-induced cardiotoxicity (76, 77). The β module, containing proteins with increased abundance due to DOX treatment, shows enrichment for both metabolic processes as well as processes involved in the translocation of enzymatic proteins to the nucleus, uniquely placing it at the intersection between the proteomic and metabolomic response to DNA damage. This includes the hub protein PRDX1, which has been shown to translocate to the nucleus upon DNA double-strand breakage and clear damage-induced nuclear reactive oxygen species and γH2AX (78). Similarly, four TCA cycle proteins help prevent DOX-mediated cellular damage by translocating from the mitochondria to the nucleus upon DNA damage (79). Three of these proteins, PDH-E1, MDH-2 and CS, are co-expressed in the β module. Therefore, the β module may identify proteins for future studies investigating how the nuclear translocation of enzymatic proteins can attenuate DOX-induced cardiotoxicity.

### Highly-connected DNA damage-associated proteins are intolerant to mutation

Our network analysis identified not only modules of co-expressed proteins but also subsets of highly connected hub proteins belonging to DOX-correlated and non-DOX-correlated modules. This allowed us to investigate the relationship between DNA damage-associated connectivity and tolerance of these proteins to physiological and pathological variation. We find that DNA damage-associated hub proteins are depleted for pQTLs, but are enriched for loss-of-function intolerant proteins indicating the importance of these proteins. Considering the spectrum of connectivity, beyond the highly-connected hub proteins, revealed increasing pLI values with increasing protein connectivity for DOX-correlated module proteins. pLI values remain low across a range of connectivities for non-DOX-correlated module proteins. The trend for DOX-correlated proteins was observable in both total connectivity and intra-modular connectivity, but not inter-modular connectivity, suggesting that intolerance to mutation in the network is centered around DOX-correlated modules and their related biological processes. The opposite trend is true for pQTLs, where enrichment tends to decrease with increased connectivity of DOX-correlated proteins. The differential relationship between connectivity, genetic influence and pLI, emphasizes the evolutionary constraint of DOX-correlated hub proteins by purifying selection to minimize potential disruptions in their expression.

### Highly-connected DNA damage-associated proteins influence CVD risk proteins through protein-protein interactions

While CVD risk proteins are not enriched amongst hub proteins or DOX-correlated proteins, physical protein interactors of CVD risk proteins are enriched amongst DOX-correlated proteins and DOX-correlated hub proteins in particular. For example, DOX-correlated hub proteins PBRM1 and SMARCC1 both interact with atrial fibrillation-associated protein DPF3, and DOX-correlated hub protein HSPA4 interacts with dilated cardiomyopathy-associated BAG3. DOX-correlated hub proteins TFB1M and TFAM interact with POLRMT in anthracycline-induced cardiotoxicity, while another risk protein, RPl7, interacts with 28 DOX-correlated hub proteins. These findings are also corroborated by the finding that GWAS risk proteins are generally very tolerant to mutation, but their physical PPIs are not. However, there are exceptions to this trend such as is observed in atrial fibrillation, where GATA4, GTF2I, and CASZ1 are both CVD proteins and DOX-correlated hub proteins. These data support the notion that genetic variation could contribute to CVD phenotypes by altering the stability and functionality of regulatory proteins that are central to the proteomic DNA damage response network through physical protein interactions. Therefore, our network pinpoints CVD risk proteins that are highly connected to proteins central to the cardiomyocyte DNA damage response that can be prioritized for cell-type specific co-immunoprecipitation studies that have proved informative for understanding mechanisms behind genes implicated in coronary artery disease and autism spectrum disorder (22, 23).

### Considering the DNA damage response network in the context of the omnigenic model

The omnigenic model for complex phenotypes posits that thousands of genes expressed in disease-relevant cell types can influence complex traits, whereby core genes have a direct impact on the phenotype, while peripheral genes influence it indirectly through distant gene interactions (80). In an omnigenic architecture, the majority of the heritability influencing complex traits with polygenic architecture is distributed across peripheral genes and involves extensive regulatory networks connecting them to core genes (80, 81). In practice, core genes are likely identified on a graded scale rather than a binary classification, where heritability decreases as the degree of separation from core genes is reduced. The use of co-expression networks to investigate the omnigenic model provides a powerful approach to untangle the complex interactions specified in this framework.

Co-expression networks across different conditions or tissues can use connectivity as a method to identify core and peripheral genes and specify the distribution of heritability across degrees of separation. For example, Mähler *et al*., used a transcriptional co-expression network from *P. tremula* leaf buds to demonstrate that eQTL genes are predominantly located at the network’s periphery, and that connectivity is inversely correlated with eQTL effect sizes, implying that core genes (hub genes) within modules experience strong selective pressures (82). The purifying selection predominantly acting on core genes implies an evolutionary conservation that possibly underscores their fundamental biological roles. This finding is also consistent with the notion that damage to core genes by loss-of-function mutations tends to have strong effects on disease risk (80, 82). Fóthi *et al*., evaluated the omnigenic model’s application to autism spectrum disorder using brain-specific gene interaction networks and found that autism gene clusters are significantly more connected to each other and the peripheral genes in brain-related tissues than in non-brain-related tissues (83). These data support the notion that disease-relevant tissues are the appropriate context for assessing omnigenic architectures to better understand complex traits. Hartl *et al*., used this foundation to generate a co-expression network derived from gene expression profiles across 12 brain regions to contextualize the functional pathways of risk genes for multiple neuropsychiatric diseases (18). Despite the omnigenic model’s suggestion that disease risk is mediated by a small number of core genes indirectly influenced by peripheral genes, the study indicated that gene effects are distributed more continuously across the networks rather than being segregated into distinct core and peripheral categories.

The omnigenic model suggests that the connections between peripheral and core genes vary by trait and include transcriptional networks, post-translational modifications, and protein-protein interactions (80, 84) and that genetic variants influencing disease may affect expression in specific cell types or conditions (80). The aforementioned studies all utilize steady-state expression data to construct their networks, whereas an omnigenic architecture relevant to disease may emerge under the conditions of specific cell stressors that drive the purifying selection pressure that core genes are placed under. The results obtained for DOX-correlated modules resemble what would be predicted by an omnigenic architecture, where there is a negative correlation between the influence of genetic variation and network connectivity. However, the opposite is true for proteins that are not correlated to DOX. This suggests that the core-periphery structure may develop in response to selective pressures. Our dynamic network differentiates proteins that are actively responding to stress from those that are not. While modules derived from steady-state networks can enrich for various biological processes, the connectivity observed at steady state may not accurately reflect gene relationships under disease-relevant conditions. Instead, the core-periphery structure may be dynamic, shifting according to different cellular states. If true, it means that networks constructed in response to various stressors might establish a connectivity profile that pinpoints unique or shared core-peripheral relationships to different cell states. Therefore, networks generated from cell models that capture the dynamic molecular responses to selective pressures or disease-associated stimuli may be needed to more accurately understand how the core-periphery structure described in the omnigenic model emerges in the transition from non-disease to disease states.

### Potential limitations of our model

We generate cardiomyocytes through directed differentiation of iPSCs as it allows us to include multiple individuals, and use these cardiomyocytes in carefully-controlled experiments where we can treat the same batch of cells with DOX and measure their protein abundance. However, it is possible that our *in vitro* system may not fully recapitulate the *in vivo* molecular profile. We also selected a single, sub-lethal, dose of DOX to study the primary effects of DNA damage on the proteome. It is possible that different doses of DOX would apply different selective pressures on the proteome and identify different response proteins and mutation tolerance. However, we believe our results using a clinically-relevant dose of a widely-used chemotherapeutic agent allow us to provide useful insights. Similarly, the response to DNA damage may be temporally dynamic and hence our results may not extrapolate to shorter or longer exposure times.

While mass spectrometry is a powerful tool for measuring protein abundance, it has inherent limitations that can affect the comprehensiveness and accuracy of the data acquired. For example, it may not detect low-abundance proteins effectively, leading to an incomplete representation of the proteome. Further, while protein abundance is a critical metric, it is not the only relevant measure when assessing protein function and cellular responses. Other important protein metrics include post-translational modifications such as phosphorylation, ubiquitination, and glycosylation, which can significantly alter protein function, stability, localization, and interactions. Future studies that integrate more comprehensive transcriptional and proteomic networks that capture multiple timepoints and greater protein coverage may enhance the findings from similarly designed studies.

In summary, there are no studies integrating proteomic data from cardiomyocytes subjected to DNA damage with measures of genetic tolerance to variation and disease. Here, we profiled protein abundance in cardiomyocytes treated with DOX across multiple individuals. We found that the level of protein connectivity in DNA damage-associated co-expression modules influences the tolerance to genetic variation. We believe that the data and analysis presented here will be a resource for further studies into the mechanistic effects of DNA damage on the cardiomyocyte proteome and DOX-induced cardiotoxicity, as well as for studies investigating the architecture of complex traits in response to perturbation.

## Methods

### Ethics statement

iPSC lines from individuals 1 and 2 (Individual 1: UCSD131i-77-1 and Individual 2: UCSD143i-87-1) were obtained from the iPSCORE resource generated by Dr. Kelly A. Frazer at the University of California San Diego as part of the National Heart, Lung and Blood Institute Next Generation Consortium (85). The iPSC lines were generated with approval from the Institutional Review Boards of the University of California San Diego and The Salk Institute (Project number: 110776ZF), and informed written consent from participants. The cell lines are available through the biorepository at WiCell Research Institute (Madison, WI, USA), or through contacting Dr. Kelly A. Frazer at the University of California, San Diego.

The iPSC line from Individual 3 (WTSIi048-A) was obtained from the HipSci project funded by the Wellcome Trust and Medical Research Council (86). The HipSci line was approved by the East of England - Cambridge Central Research Ethics Committee (REC 15/EE/0049). The cell line was generated with informed consent of the participant. The cell line is available from the European Bank of induced pluripotent Stem Cells (EBiSC) and European Collection of Authenticated Cell Cultures (ECACC).

### Induced pluripotent stem cell lines

The iPSCORE iPSC lines: UCSD143i-87-1 and UCSD131i-77-1 were generated from skin fibroblasts from two unrelated, healthy female donors of Asian-Chinese ethnicity aged 23 and 21 respectively. The HipSci iPSC line: WTSIi048-A was generated from skin fibroblasts from a 72-year-old female of White British ethnicity. All donors used in our study were healthy with no previous history of cardiac disease. All lines tested negative for mycoplasma contamination.

### iPSC culture

Feeder-independent iPSCs were cultured in mTESR1 (85850, Stem Cell Technology, Vancouver, BC, Canada) media with 1% Penicillin/Streptomycin (30-002-Cl, Corning, Bedford, MA. USA) on hESC-qualified Matrigel Matrix (354277, Corning, Bedford, MA, USA) at a dilution of 1:100. iPSCs were passaged with dissociation reagent (0.5 mM EDTA, 300 mm NaCl in PBS) when they attained 70-80% confluency, approximately every 3-5 days.

### Cardiomyocyte differentiation

Cardiomyocyte differentiations were performed as previously described (26). Briefly, iPSC lines were seeded in Matrigel-coated culture dishes (Days -6/-5) and cultured until 85–90% confluent. Differentiations were initiated (Day 0) by adding 12 μM of the GSK3 inhibitor, CHIR99021 trihydrochloride (4953, Tocris Bioscience, Bristol, UK) in Cardiomyocyte Differentiation Media (CDM) [500 mL RPMI 1640 (15-040-CM, Corning), 10 mL B-27 minus insulin (A1895601, ThermoFisher Scientific, Waltham, MA, USA), 5 mL GlutaMAX (35050-061, ThermoFisher Scientific), and 5 mL of Penicillin/Streptomycin (100X) (30-002-Cl, Corning)]. After 24 hr (Day 1), the media was replaced with fresh CDM. On Day 3 (after 48 hr) media was replaced with CDM containing 2 μM of the Wnt signaling inhibitor Wnt-C59 (5148, Tocris Bioscience) in CDM. CDM was replaced on Day 5, 7, 10 and 12. We observed spontaneously beating cells between Day 7–10. iPSC-CMs were purified by metabolic selection with glucose-free, lactate-containing media [500 mL RPMI without glucose (11879, ThermoFisher Scientific), 106.5 mg L-Ascorbic acid 2-phosphate sesquimagnesium salt (sc228390, Santa Cruz Biotechnology, Santa Cruz, CA, USA), 3.33 ml 75 mg/ml Human Recombinant Albumin (A0237, Sigma-Aldrich, St Louis, MO, USA), 2.5 mL 1 M lactate in 1 M HEPES (L(+)Lactic acid sodium (L7022, Sigma-Aldrich)), and 5 ml Penicillin/Streptomycin] added on Day 14, 16 and 18. On Day 20, iPSC-CMs were detached with 0.05% Trypsin-EDTA solution (25–053 Cl, Corning), and a single cell suspension was generated by straining. iPSC-CMs were counted with a Countess 2 machine. 1.5 million iPSC-CMs were plated per well of a 0.1% Gelatin-coated 6-well plate in 3 mL Cardiomyocyte Maintenance Media (CMM) [500 mL DMEM without glucose (A1443001, ThermoFisher Scientific), 50 mL FBS (S1200-500, Genemate), 990 mg Galactose (G5388, Sigma-Aldrich), 5 mL 100 mM sodium pyruvate (11360–070, ThermoFisher Scientific), 2.5 mL 1 M HEPES (SH3023701, ThermoFisher Scientific), 5 mL Glutamax (35050–061, ThermoFisher Scientific), 5 mL Penicillin/Streptomycin]. The iPSC-CMs were matured in culture for 10 days, with CMM replaced on Day 23, 25, 27, 28, and 30.

### iPSC-CM purity determination using flow cytometry

After differentiation of iPSCs from Individuals 2 & 3, the iPSC-CM purity was determined using flow cytometry. Between Day 25-27 of differentiation the iPSC-CMs were dissociated with 0.05% Trypsin-EDTA solution and strained to generate a single cell suspension. One million cells were stained with Zombie Violet Fixable Viability Kit (423113, BioLegend) for 30 min at 4 °C prior to fixation and permeabilization (FOXP3/Transcription Factor Staining Buffer Set, 00– 5523, ThermoFisher Scientific) for 30 min at 4 °C. Cells were stained with 5 ml PE Mouse Anti-Cardiac Troponin T antibody (564767, clone 13–11, BD Biosciences, San Jose, CA, USA) for 45 min at 4 °C. Cells were washed three times in permeabilization buffer and re-suspended in autoMACS Running Buffer (130-091-221, Miltenyi Biotec, Bergisch Gladbach, Germany). We used several negative controls in each flow cytometry experiment: 1) iPSCs, which should not express TNNT2, 2) an iPSC-CM sample that has not been labeled with viability stain or TNNT2 antibody, 3) an iPSC-CM sample that is only labeled with the viability stain and 4) an iPSC-CM sample that is only labeled with TNNT2. 10,000 cells were captured and profiled on the BD LSRFortessa Cell Analyzer. Multiple gating steps were performed to determine the proportion of TNNT2-positive cells: 1) Cellular debris was removed by gating out cells with low granularity on FSC versus SSC density plots, 2) From this population, live cells were identified as the violet laser-excitable, Pacific Blue dye-negative population, 3) TNNT2-positive cells were identified within the set of live cells and any cells that overlap the profiles of the negative control samples were excluded. iPSC-CM purity is reported as the proportion of TNNT2-positive live cells.

### Drug treatment of iPSC-CMs

On Day 29, iPSC-CMs were treated with 0.5 μM of Doxorubicin (D1515, Sigma-Aldrich) or vehicle (Molecular Biology grade water) in fresh CMM media for 24 hr. The treatment for Individual 3 was replicated two additional times yielding 10 samples in total across three individuals. Post-treatment, cells were washed twice and scraped in ice-cold PBS. iPSC-CMs were flash-frozen and stored at -80 °C prior to further processing.

### γH2AX immunofluorescence staining and quantification of DNA double-strand breaks

300,000 iPSC-CMs were seeded per well of a 24-well plate in CMM media. Cells were treated with 0.5 μM DOX or vehicle (DMSO). The treated cells were fixed in 4% paraformaldehyde for 15 min and permeabilized with 0.25% DPBS-T for 10 min at room temperature. Cells were incubated with 5% BSA:DPBS-T for 30 min at room temperature, then incubated with a 1:500 dilution of γ-H2AX primary antibody in BSA:DPBS-T overnight at 4 °C (Phospho-Histone H2A.X (Ser139) Rabbit mAb; NC1602516; Fisher Scientific). Cells were incubated with Fluorochrome-conjugated secondary antibody (Donkey anti-Rabbit Alexa Fluor 594, A-21207, Invitrogen) for 1 hr at room temperature at a 1:1000 dilution in DPBS-T. Cell nuclei were counterstained with Hoechst 33342 nucleic acid stain (PI62249, Thermo Scientific) for 10 min in the dark. Stained cells were subjected to fluorescence microscopy. The total number of nuclei and γH2AX-positive nuclei were quantified using the cell counter plugin of ImageJ software (87). The number of γH2AX-positive nuclei were divided by the total number of nuclei to determine the percentage of γH2AX-positive nuclei in DOX- and vehicle-treated iPSC-CMs for three different individuals. The percentage of γH2AX-positive cells between vehicle- and DOX-treated iPSC-CM samples was compared by t-test.

### Protein isolation and quantification

Protein was isolated from iPSC-CMs by lysing the cells with RIPA buffer [1.5 ml 5 M NaCl, 1 ml Triton X100, 1 g Na deoxycholate, 1 ml 10% SDS, 1 ml 1 M Tris pH 7.4 and 45 ml dH_2_O] with protease inhibitor for 1 hr at 4 °C. Isolated proteins were quantified by using the BCA Protein Assay kit (23227, Thermo Scientific) according to the manufacturer’s instructions.

### Protein digestion

The samples were prepared similarly as previously described (88). Briefly, 15 μg of protein were solubilized with 60 μL of 50 mM Triethylammonium bicarbonate (TEAB) pH 7.55. The proteins were then reduced with 10 mM Tris(2-carboxyethyl) phosphine (TCEP) (77720, Thermo) and incubated at 65 °C for 10 min. The sample was then cooled to room temperature and 1 μL of 500 mM iodoacetamide acid was added and allowed to react for 30 min in the dark. Then, 3.3 μl of 12% phosphoric acid was added to the protein solution followed by 200 μL of binding buffer (90% Methanol, 100mM TEAB pH 8.5). The resulting solution was added to S-Trap spin column (protifi.com) and passed through the column using a bench top centrifuge (60 s spin at 1,000 g). The spin column is washed with 150 μL of binding buffer and centrifuged. This is repeated two times. 30 μL of 20 ng/μL Trypsin is added to the protein mixture in 50 mM TEAB pH 8.5, and incubated at 37 ^○^C overnight. Peptides were eluted twice with 75 μL of 50% acetonitrile, 0.1% formic acid. Aliquots of 20 μL of eluted peptides were quantified using the Quantitative Fluorometric Peptide Assay (Pierce, Thermo Fisher Scientific). Eluted volume of peptides corresponding to 5.5 μg of peptides are dried in a speed vac and resuspended in 27.5 μL 1.67% acetonitrile, 0.08% formic acid, 0.83% acetic acid, 97.42% water and placed in an autosampler vial.

### Nanoflow liquid chromatography mass spectrometry

Peptide mixtures were analyzed by nanoflow liquid chromatography-tandem mass spectrometry (nanoLC-MS/MS) using a nano-LC chromatography system (UltiMate 3000 RSLCnano, Dionex), coupled on-line to a Thermo Orbitrap Eclipse mass spectrometer (Thermo Fisher Scientific, San Jose, CA) through a nanospray ion source. Instrument performance was verified by analyzing a standard six protein mix digest before the sample set run, between each experimental block and at the end of the experiment. The six protein mix data files were analyzed to confirm that instrument performance remained consistent throughout the experiment. A direct injection method using 3 μL of digest onto an analytical column was used; Aurora (75 µm X 25 cm, 1.6 µm) from (IonOpticks). After equilibrating the column in 98% solvent A (0.1% formic acid in water) and 2% solvent B (0.1% formic acid in acetonitrile (ACN)), the samples (2 µL in solvent A) were injected (300 nL/min) by gradient elution onto the C18 column as follows: isocratic at 2% B, 0-5 min; 2% to 6%, 5-5.1 min; 6% to 25% 5.1-105 min, 25% to 50% B, 105-120 min; 50% to 90% B, 120-122 min; isocratic at 90% B, 122-124 min; 90% to 5%, 124-125 min; isocratic at 5% B, 125-126 min; 5% to 90% 126-128 min; isocratic for one min; 90%-2%, 129-130 min; and isocratic at 2% B, till 150 min.

### NanoLC MS/MS Analysis for DDA

All data were acquired using an Orbitrap Eclipse in positive ion mode using a top speed data-dependent acquisition (DDA) method with a 3 s cycle time and a spray voltage of 1600 V. The survey scans (m/z 375-2000) were acquired in the Orbitrap at 120,000 resolution (at m/z = 400) in profile mode, with a maximum injection time of 100 ms and an AGC target of 1,000,000 ions. The S-lens RF level was set to 30. Isolation was performed in the quadrupole with a 1.6 Da isolation window, and HCD MS/MS acquisition was performed in profile mode using the orbitrap at a resolution of 15,000 using the following settings: parent threshold = 5,000; collision energy = 30%; AGC target at 125,000 using the default settings. Monoisotopic precursor selection (MIPS) and charge state filtering were on, with charge states 2-10 included. Dynamic exclusion was used to remove selected precursor ions, with a +/- 10 ppm mass tolerance, for 30 s after acquisition of one MS/MS spectrum.

### NanoLC MS/MS Analysis for DIA

All LC-MS/MS data were acquired using an Orbitrap Eclipse in positive ion mode using a data-independent acquisition (DIA) method with 16 Da windows from 400-1000 and a loop time of 3 s. The survey scans (m/z 350-1500) were acquired in the Orbitrap at 60,000 resolution (at m/z = 400) in centroid mode, with a maximum injection time of 118 ms and an AGC target of 100,000 ions. The S-lens RF level was set to 60. Isolation was performed in the quadrupole, and HCD MS/MS acquisition was performed in profile mode using the orbitrap at a resolution of 30000 using the following settings: collision energy = 33%, IT 54 ms, AGC target = 50,000. A pooled sample was used to create spectral libraries that we search the individual samples against by injecting 5 times using narrow (4 Da), staggered windows over 100 m/z ranges from 400-900 m/z in a technique called gas phase fractionation as described in Searle *et al*. (89).

### Database searching for DDA proteins

Tandem mass spectra were extracted and charge state deconvoluted using Proteome Discoverer (Thermo Fisher, version 2.2.0388). Deisotoping was not performed. All MS/MS spectra were searched against a Uniprot Human database using Sequest and the Minora node used to perform Label-Free Quan (LFQ) using the MS peak areas for each of the peptide-spectral matches (PSMs). Searches were performed with a parent ion tolerance of 5 ppm and a fragment ion tolerance of 0.02 Da. Trypsin was specified as the enzyme, allowing for two missed cleavages. Fixed modification of carbamidomethyl (C) and variable modifications of oxidation (M) and deamidation were specified in Sequest. Protein identities reported at 1% false discovery rate were considered for filtering and downstream analysis.

### Database searching for DIA proteins

The raw data was demultiplexed to mzML with 10 ppm accuracy after peak picking in MSConvert (90). The resulting mzML files were searched in MSFragger (91) and quantified via DIA-NN (https://github.com/vdemichev/DiaNN) using the following settings: peptide length range 7-50, protease set to Trypsin, 2 missed cleavages, 3 variable modifications, clip N-term M on, fixed C carbamidomethylation, variable modifications of methionine oxidation and n-terminal acetylation, MS1 and MS2 accuracy set to 20 ppm, 1% FDR, and DIANN quantification strategy set to Robust LC (high accuracy). The files were searched against a database of human acquired from Uniprot (18^th^ December, 2023). The gas-phase fractions were used only to generate the spectral library, which was used for analysis of the individual samples.

### Abundance matrix filtering and imputation

We removed four non-human or uncharacterized proteins (UniProt: Q6ZSR9, P15252, P25691, P00761). Proteins that were missing across 50% or more of the samples were removed. For the remaining 246 proteins, missing values were imputed using k-Nearest Neighbors from the impute package knn. with, parameters k (number of neighbors to be used in the imputation) and rowmax (the maximum percent missing data allowed in any row) (30). Using k = 10 in the impute.knn function leverages all available samples to impute missing values, given the small sample size of our dataset. The rowmax = 0.4 parameter allows for imputing rows with up to 40% missing values, balancing the need to retain as much data as possible while maintaining the quality of the imputations. These steps led to a total of 4,178 analyzable proteins across all samples.

### Removal of unwanted technical variation and normalization

To eliminate unwanted technical variation from the log_2_-transformed imputed abundance matrix, we adjusted the 10 sample data matrix using both negative controls and the replicate data with the RUV-III function in the ruv R package (92). Negative control proteins were defined as the 5% least variable proteins across all samples. The RUV-corrected abundance matrix and RUV factors were used in downstream analysis.

### Comparison of the iPSC-CM proteome with the proteome across human tissues

Protein abundance values across 26 tissues was obtained from Jiang *et. al* (20). We identified the set of proteins expressed across the 26 tissues and our iPSC-CMs. We calculated the median abundance across individuals for the set of 26 tissues, as well as for our iPSC-CMs. The median abundance between our iPSC-CMs and each of the 26 tissues was correlated using Pearson correlation.

### WGCNA network construction

We adopted the Weighted Gene Co-expression Network Analysis (WGCNA) methodology to investigate correlations between protein abundances in our RUV-corrected abundance matrix of DOX- and VEH-treated iPSC-CMs (32). The WGCNA framework, primarily utilizing wrapper functions from BioNERO, was implemented for this analysis (93). We first established a scale-free network topology, achieved by determining the appropriate soft threshold power using the SFT_fit function. After iterating different soft power thresholds (β), the linear regression of log_10_(k) versus log_10_(p(k)) indicates that by setting β = 20, the network is close to a scale-free network, where k is the whole network connectivity and p(k) is the corresponding frequency distribution. This setting resulted in our network attaining a scale-free fit index of 0.71 (quantification of how well the network approximates a scale-free topology), with a mean node connectivity of 65.4 and median connectivity of 53.6. We identified modules of co-expressed proteins using a modified workflow as previously described (32, 93). The workflow was encapsulated in the exp2gcn2 function from the BioNERO package. We selected a signed network with a soft power threshold of 20, merging threshold of 0.85, and the pearson correlation method. We computed the Topological Overlap Matrix (TOM), a measure of network connectivity that emphasizes the shared neighbors between protein pairs to enhance the robustness and reliability of the calculated adjacency network. Using hierarchical clustering on the dissimilarity TOM (dissTOM), we identified initial modules of co-expressed proteins. The dynamic tree cut method using the cutreeDynamicTree function with maxTreeHeight of 3, minimum module sizes of 40 and no deep splitting was applied to the protein dendrogram to define these modules by modifying the exp2gcn2 function. The module eigenproteins (MEs), representing the first principal component for the module, were then calculated across modules with moduleEigengenes. These eigenproteins served as characteristic expression profiles of proteins within a module and were used to assess the interrelation between modules. Similar modules were merged based on the eigenprotein dissimilarity, ensuring that highly correlated modules were combined. Modules whose eigenproteins had a correlation of 0.85 or greater were merged.

We used three types of connectivity metrics to describe how each node/protein in the network related to other nodes/proteins. Total connectivity was calculated by summing the weighted correlations between each protein and all other proteins in the network (kTotal). Intra-modular connectivity (kIN) was calculated by summing the weighted correlations between each protein and all other proteins in the assigned module. Extra-modular connectivity (kOut) was calculated by summing the weighted correlations between each protein and all other proteins outside the assigned module.

### Identification of hub proteins

We identified hub proteins that might play central roles in the biological processes represented by each module. We used the get_hubs_gcn function to identify hub proteins within our protein abundance correlation network. Hub proteins within modules are defined as the proteins with the highest intra-modular connectivity (kIN) score (top 10%) and the highest pearson correlation value with the module eigenprotein (> 0.8).

### Module correlation to DOX treatment and individual

To assess the relationship between each module’s eigenprotein and the defined trait (DOX treatment or individual), we performed a pearson correlation analysis using the cor.test function from the stats package in R (94). This analysis provided a correlation coefficient for each module, indicating the strength and direction of the association between the module’s expression profile and the experimental condition. The statistical significance of these correlations was determined by extracting *P* values for each module-trait correlation using the cor.test function, where *P* values less than 0.01 were determined to be significant.

### Hub protein correlation network visualization in DOX-correlated modules

The correlation network for hub proteins in DOX-correlated modules was visualized using the igraph package in R (95). Abundance correlations between proteins were obtained using the get_edge_list function from BioNERO (93). Protein pairs with positive Pearson correlations of 0.9 or more were selected for visualization.

### Linear modelling to identify differentially abundant proteins

We utilized the limma package to fit protein abundances to a linear model across conditions (96, 97). We randomly selected one technical replicate per treatment group from Individual 3 so as not to confound technical and biological variation in the linear modelling process. The RUV-III corrected abundance matrix from six samples was quantile normalized using the normalizeBetweenArrays function. Drug treatment (DOX or VEH) was modelled as a fixed effect, whereas individual (IND) was treated as a random effect estimated using the duplicateCorrelation function. The linear model fitting was done using lmFit, which incorporated the block effect from the individuals and the design matrix. This model was then passed through the empirical Bayes moderation in the eBayes function to obtain moderated t-statistics. We defined contrasts in the linear model to compare the differential expression between DOX and VEH conditions such that positive log_2_-fold change values correspond to increased abundance in the DOX-treated group, and negative values correspond to decreased abundance in the DOX-treated group. The model was refitted with these contrasts, and empirical Bayes moderation was applied again to adjust the statistics. This summary was visually explored through a histogram of nominal *P* values, whereby observation of the distribution and plotting of abundance values across samples led us to determine that a nominal *P* < 0.05 was an appropriate cutoff for significance for differential abundance. We denote proteins that meet this criterion as differentially abundant proteins (DAP).

### Comparison of the DIA and DDA datasets

We detected 4,501 proteins by DDA and removed all proteins containing at least one missing protein abundance value, resulting in 3,384 measured proteins. We detected 4,261 proteins by DIA and removed all proteins containing at least one missing protein abundance value, resulting in 3,934 measured proteins. We selected the 3,027 proteins shared between these two sets and calculated the mean log_2_ abundance for each protein across all samples. DDA and DIA protein log_2_ abundance were fit to a linear model using the lmFit function to determine the strength and significance of the correlation between data sets. We identified differentially abundant proteins in the DDA data set as described above for DIA, using the same preprocessing and modelling parameters. We then compared the response effect sizes (log_2_ fold change) between proteins shared between the two data sets using the lmFit function to determine the strength and significance of the correlation between data sets.

### Comparison of proteins elevated in heart tissue across modules

We sourced data on proteins whose expression is elevated in heart tissue compared to other tissue types from the Human Protein Atlas (33). Elevated proteins correspond to those with at least a four-fold higher mRNA level in a particular tissue compared to any other tissue. Proteins identified as elevated in heart tissue were categorized into their respective modules within our correlation network. We then calculated their percentages relative to the total protein count within each respective module.

### Comparison of heart ventricle tissue specificity scores across modules

Tissue specificity values for ‘Heart.Ventricle’ were obtained for our 4,178 proteins from Supplementary Table 2 of (20). We tested for differences in tissue specificity across modules using a Wilcoxon rank-sum test for each module. We compared scores between all proteins in a given module, and all proteins not contained within that module, where significant differences were denoted when *P* < 0.05. The same criteria and test were also applied to compare tissue specificity between DAPs and non-DAPs.

### Cellular localization of module proteins

We retrieved lists of proteins classified as signal peptides, voltage-gated channels, secreted, intracellular, membrane-bound, or plasma proteins from the Human Protein Atlas database (33). We utilized the UniProt database (34) to identify proteins experimentally shown to be located in subcellular structures including autophagosomes, membranes, cytoplasm, cytoskeleton, endoplasmic reticulum, Golgi apparatus, lysosomes, mitochondria, nucleus, and sarcomere. We classified each protein in the correlation network according to the above annotations, and performed a Fisher’s exact test to determine which modules were enriched for proteins from a particular classification. Fisher’s exact test *P* < 0.05 was considered significant.

### Biological process enrichment across modules

To interpret the biological significance of the WGCNA-derived modules correlated with the DOX treatment, enrichment analysis was performed based on annotated Gene Ontologies (GO) (98). The enrichGO function from the clusterProfiler R package was used to test the enrichment of terms associated with biological processes against the background of all network proteins (99). Enriched terms were those with a Benjamini-Hochberg adjusted *P* < 0.05.

### Functional categorization of module proteins

We retrieved lists of proteins classified as human transcription factors (35), RNA binding proteins (RBPs) (37), enzymes and transporter proteins (33), and mammalian stress granule (MSG) proteins (36). Enrichment of different protein categories within each module was determined by Fisher’s exact test.

### Protein family enrichment amongst module proteins

We queried UniProt for protein family names for the set of network proteins (34). We assigned each protein to its set of families and performed a Fisher’s exact test to determine which modules were enriched for proteins from a particular family. Fisher’s exact *P* < 0.05 was considered significant.

### pQTL protein enrichment

We obtained plasma pQTL data from the UK Biobank from Supplementary Table 10 (17). We selected pQTLs that were identified independent of genetic ancestry. pQTLs were further classified as *cis*-pQTLs or *trans*-pQTLs. For pQTL protein enrichment analysis, we collated the set of unique pQTL proteins. We tested for enrichment of pQTL proteins among hub proteins, DOX-correlated proteins, and DOX-correlated hub proteins, compared to the set of proteins not contained within each of those sets using the Fisher’s exact test. A Fisher’s exact *P* < 0.05 was considered statistically significant.

### Comparison of pQTL SNP effect sizes

From the aforementioned set of *cis*-pQTLs and *trans*-pQTLs, we selected the SNP with the highest effect size for each protein. We tested for differences in effect sizes between DOX-correlated and non-DOX-correlated proteins, hub proteins and non-hub proteins, and DOX-correlated hub proteins and non-DOX-correlated hub proteins using a Wilcoxon rank-sum test, where a *P* < 0.05 was considered statistically significant.

### Proportion of pQTLs across network protein connectivity deciles

For each protein in the network, we calculated the normalized intra-modular connectivity by taking the kIN of each protein and dividing it by one less than the number of module connections. Proteins were assigned to one of two groups based on their DOX-correlation status, and stratified into deciles based on their normalized intra-modular connectivity. We then calculated the proportion of proteins in each decile that were pQTLs (either *cis-* or *trans*-pQTLs).

### pLI, pHaplo and pTriplo comparisons across modules

We obtained probability of loss-of-function intolerance (pLI) scores (100) and probability of Haploinsufficiency (pHaplo) and Triplosensitivity (pTriplo) scores (43). We tested whether the set of scores for each metric was different for each module compared to all proteins in the network outside the module using the Wilcoxon rank-sum test. A *P* < 0.05 was considered to be significant.

### Comparisons between pLI scores and network connectivity

Proteins with a pLI ≥ 0.9 are considered intolerant to mutation, while proteins with a pLI ≤ 0.1 are considered tolerant to mutation. We compared the normalized kIN values of mutation-intolerant proteins to mutation-tolerant proteins using the Wilcoxon rank-sum test. A *P* < 0.05 was considered to be significant.

We used three types of connectivity metrics to describe how each protein in the network related to pLI scores across connectivity deciles. Total connectivity was calculated by summing the weighted correlations between each protein and all other proteins in the network (kTotal). Normalized kIN was used as described above. Extra-modular connectivity (kOut), the correlation between proteins within each module to all proteins outside the module, was normalized by dividing kOut by the number proteins outside the target protein’s module. Proteins were assigned into two groups based on their DOX-correlation status, and deciles generated for each aforementioned type of connectivity.

To ascertain enrichment for mutation-tolerant or intolerant proteins among hub proteins, DOX-correlated proteins, and DOX-correlated hub proteins, the Fisher’s exact test was conducted. A Fisher’s exact *P* < 0.05 was considered statistically significant.

### CVD GWAS enrichment testing

We obtained GWAS summary statistics from the GWAS catalog (45). We downloaded summary statistics for “heart disease” (EFO_0003777) and “heart function measurement” (EFO_0004311). We first combined the set of mapped genes of traits identified through multi-trait analysis of GWAS (MTAG) with their non-MTAG terms to limit trait redundancy. We then combined the mapped genes of traits from “Anthracycline-induced cardiotoxicity in early breast cancer”, “Anthracycline-induced cardiotoxicity in childhood cancer”, and “Anthracycline-induced cardiotoxicity in breast cancer” into one trait termed “Anthracycline-induced cardiotoxicity” to capture the broadest set of genes for this DOX-related trait. Beginning with 1,564 traits, we filtered out 487 traits contained within the heart disease and heart function measurement summary statistics where at least one risk locus was not mapped to a gene or expressed within our network. The remaining mapped genes for each of the 1,077 traits were overlapped with those expressed as proteins in our data.

Module-wise enrichment testing was then performed by comparing to the set of all proteins not contained within the module of interest. This analysis involved constructing contingency tables for each module-gene set pair and conducting Fisher’s exact tests to determine the statistical significance of the overlap. Enriched traits are defined as those with a Fisher’s exact test *P* < 0.05 and at least two overlapped genes with module proteins.

To ascertain enrichment for CVD risk proteins among hub proteins, DOX-correlated proteins, and DOX-correlated hub proteins, the Fisher’s exact test was conducted. A Fisher’s exact *P* < 0.05 was considered statistically significant.

### CVD risk protein interactor enrichment testing

Using the set of CVD risk proteins as mentioned above, protein UniProt IDs were uploaded to STRINGdb (https://string-db.org/) and PPI networks were generated (101). We focused on proteins with experimental interaction evidence (confidence score ≥ 0.9). The protein interactors for CVD risk proteins were denoted “CVD protein interactors”. To ascertain enrichment for CVD proteins interactors among hub proteins, DOX-correlated proteins, and DOX-correlated hub proteins, the Fisher’s exact test was conducted. A Fisher’s exact *P* < 0.05 was considered statistically significant.

### pLI comparison between CVD proteins and CVD protein interactors

Using the set of CVD proteins and CVD protein interactors as mentioned above, we assigned each protein their pLI score as previously described. We compared the pLI values between the two groups using the Wilcoxon rank-sum test. A *P* < 0.05 was considered to be significant.

We also generated a degree-randomized PPI network with resampling to ensure robustness of results. We utilized the set of CVD proteins and their physical protein-protein interaction partners, as determined by STRINGdb with confidence scores above 0.9. The degree of each GWAS protein within the physical PPI network was calculated, resulting in a degree distribution for the CVD proteins. This distribution highlighted the percentage of proteins with varying degrees of interaction. CVD proteins were assigned group names based on their degree, and we subsequently calculated the degree for every protein in the entire network. Proteins in the network with degrees similar to those of the CVD proteins were assigned to the same group, while those with degrees not captured by the CVD-proteins were assigned to the group with the nearest degree. Each protein in the network was thus assigned to a group reflective of the degree distribution of the GWAS proteins, ensuring comprehensive sampling without excluding proteins based on their degree. We resampled while maintaining a degree-based proportion comparable to that of the original CVD proteins. We simulated 10,000 subnetworks among sampled proteins and their interactors, comparing pLI scores between CVD proteins and their physical interactors using the Wilcoxon rank-sum test.

### PPI network construction

UniProt IDs for all proteins in the iPSC-CM network were imported into Cytoscape (Version 3.10.2) to generate a PPI network. PPIs were determined using the STRINGdb application in Cytoscape. We selected the ‘physical subnetwork’ of PPIs with confidence scores > 0.4, prior to later sub-setting PPIs with confidence scores > 0.9 in R. The combination of both moderate and stringent cutoffs was used for sensitivity analysis of our results. We then annotated proteins in the network by whether they were DOX-correlated, hub proteins or CVD proteins, as well as by which module each protein belonged to. Edges in the network represent the WGCNA-derived weighted correlation between proteins. We next generated a subnetwork from the total network by selecting PPIs where at least one protein in the PPI was a CVD protein to center our analyses on the differences between CVD proteins and their direct physical protein interactors. We then used the tidygraph (102) and ggraph (103) packages in the R programming environment to visualize a network of CVD proteins and their physical protein interactors.

## Data availability

Mass Spectrometry data will be available from the MassIVE database (https://massive.ucsd.edu/ProteoSAFe/static/massive.jsp). All custom analysis scripts used for this project are available at (https://omar-johnson.github.io/DOX_24_Github/index.html) and generated using workflowr (104).

## Acknowledgements

We thank all members of the Ward Lab for helpful discussions. We thank Dr. Kelly A. Frazer and the University of California San Diego for providing the iPSC lines through the iPSCORE resource. We acknowledge the Wellcome Trust Sanger Institute as the source of the WTSIi048-A human induced pluripotent cell line which was generated under the Human Induced Pluripotent Stem Cell Initiative funded by a grant from the Wellcome Trust and Medical Research Council, supported by the Wellcome Trust (WT098051) and the NIHR/Wellcome Trust Clinical Research Facility, and acknowledge Life Science Technologies Corporation as the provider of Cytotune. We thank the Mass Spectrometry Facility at the University of Texas Medical Branch for performing the Mass Spectrometry experiments, and the Flow Cytometry and Cell Sorting Core Facility for access to flow cytometers. This work was funded by a Cancer Prevention Research Institute of Texas (https://www.cprit.state.tx.us/) Recruitment of First-Time Faculty Award (RR190110) to M.C.W, the National Institutes of Health grant R35GM150459 to M.C.W, and CPRIT grant RP190682 to W.K.R in partial support of the UTMB Mass Spectrometry Facility. O.D.J was supported by a Jeane B. Kempner Predoctoral Fellowship administered through UTMB (http://www.kempnerfund.org/).

## Author contributions

M.C.W conceived and designed the study. S.P, J.A.G and W.K.R performed the experiments and generated the data. O.D.J, S.P and M.C.W analyzed the data. O.D.J and S.P visualized the data. O.D.J and M.C.W wrote the paper with input from all authors. M.C.W supervised the project. All authors read and approved the final manuscript.

## Supporting information

S1 Appendix: Raw protein abundance matrix.

S2 Appendix: WGCNA-derived weighted correlations of protein abundance across all pairs of proteins in the iPSC-CM DOX response network.

S3 Appendix: Document containing Supplemental Figures 1-12. S4 Appendix: Document containing Tables 1-4.

