## Appendix S3 for "DNA damage-associated protein co-expression network in cardiomyocytes informs on tolerance to genetic variation and disease"

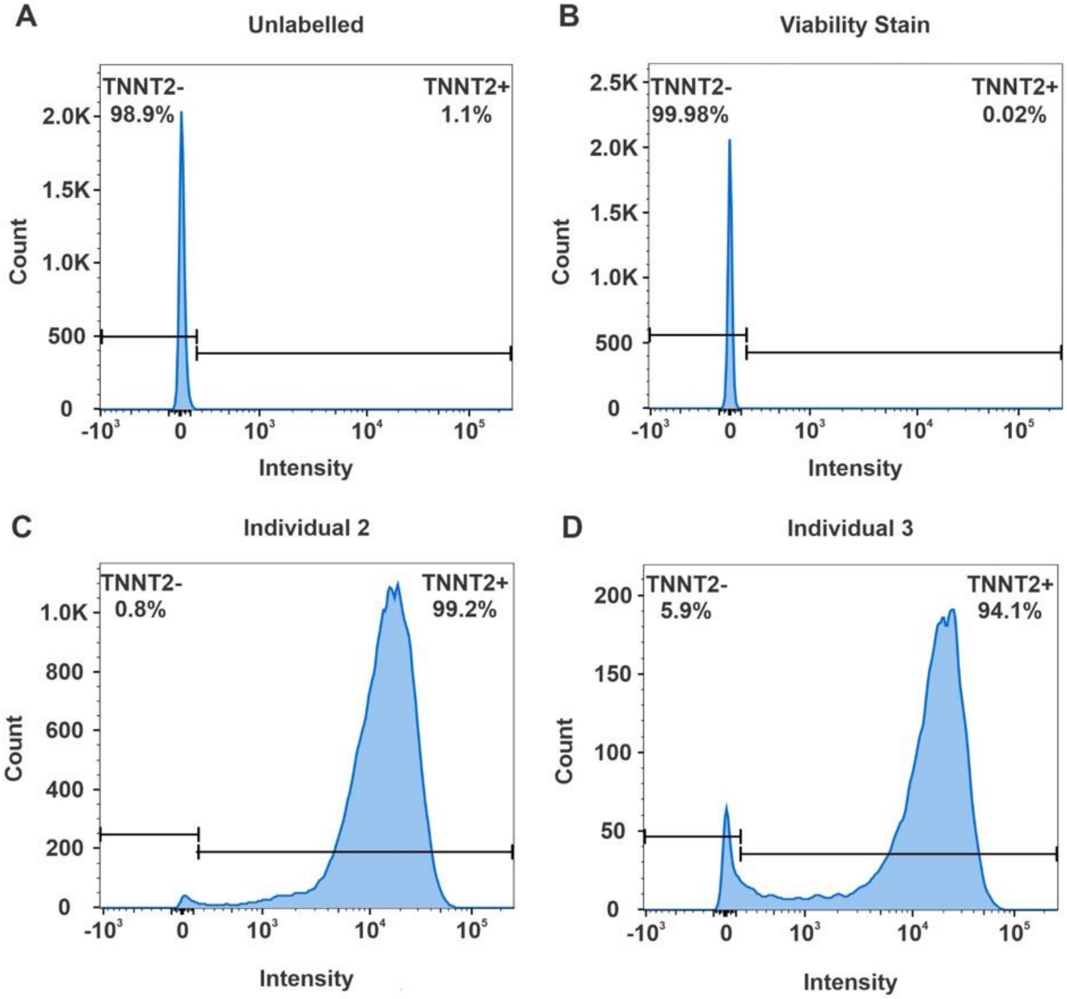


**Figure S1: iPSC-derived cardiomyocytes can be generated at high purity. (A)** Proportion of troponin (TNNT2) positive cells in unlabeled iPSC-CMs determined by flow cytometry. **(B)** Proportion of TNNT2-positive cells in iPSC-CMs labeled with viability stain. **(C)** Proportion of live TNNT2-positive cells in Individual 2 iPSC-CMs. **(D)** Proportion of live TNNT2-positive cells in Individual 3 iPSC-CMs.

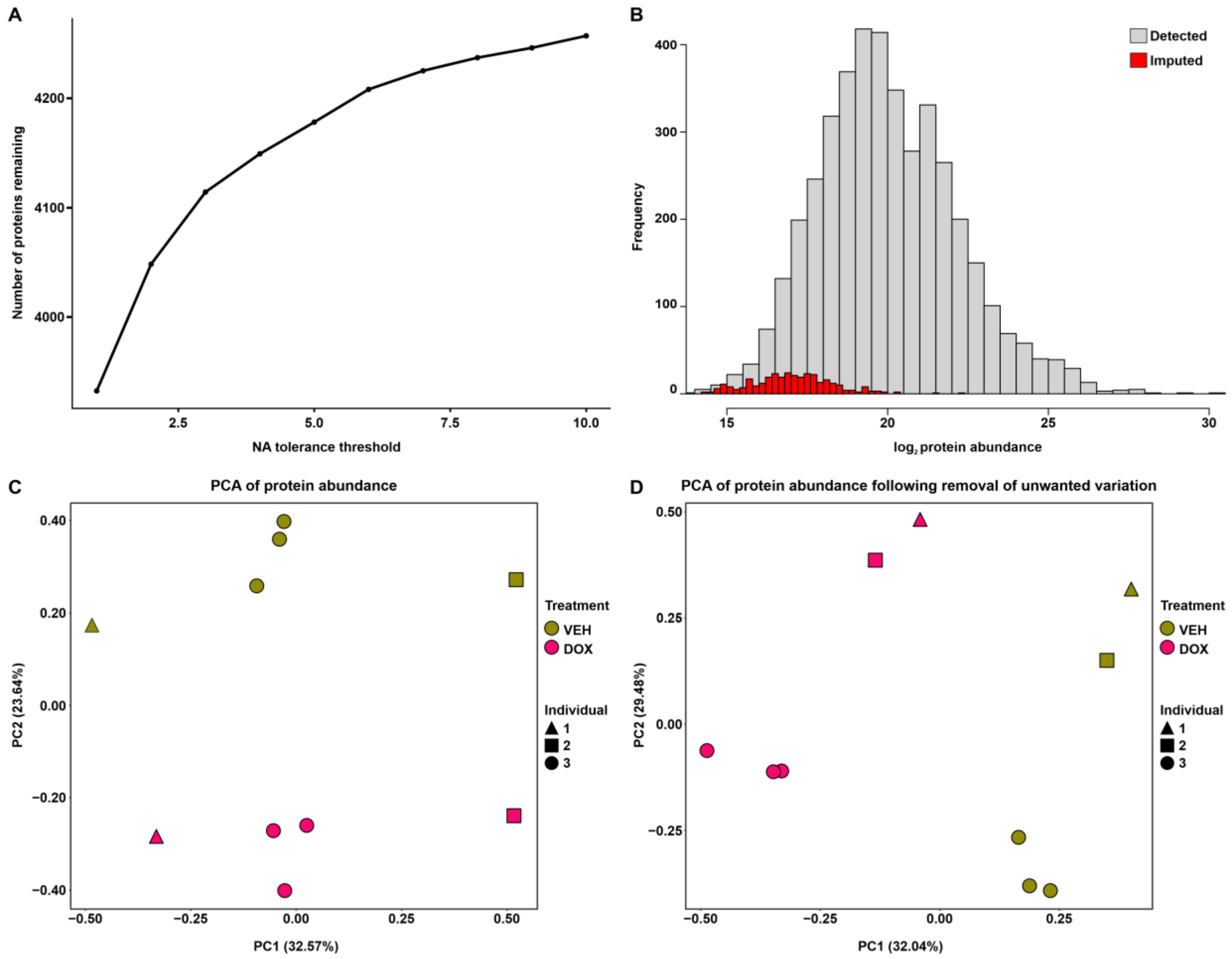


**Figure S2: DOX treatment is the primary contributor to variation in protein abundance following pre-processing. (A)** The number of measured proteins at different sample thresholds of missing values. **(B)** Distribution of detected (grey) and imputed (red) $\log_2$ protein abundance values. **(C)** Principal component analysis (PCA) of $\log_2$ protein abundance data. Samples are colored by treatment (DOX: pink, VEH: olive) and shaped by individual (Individual 1: triangle, Individual 2: square, Individual 3: circle). **(D)** PCA of $\log_2$ protein abundance values after the removal of unwanted technical variation.

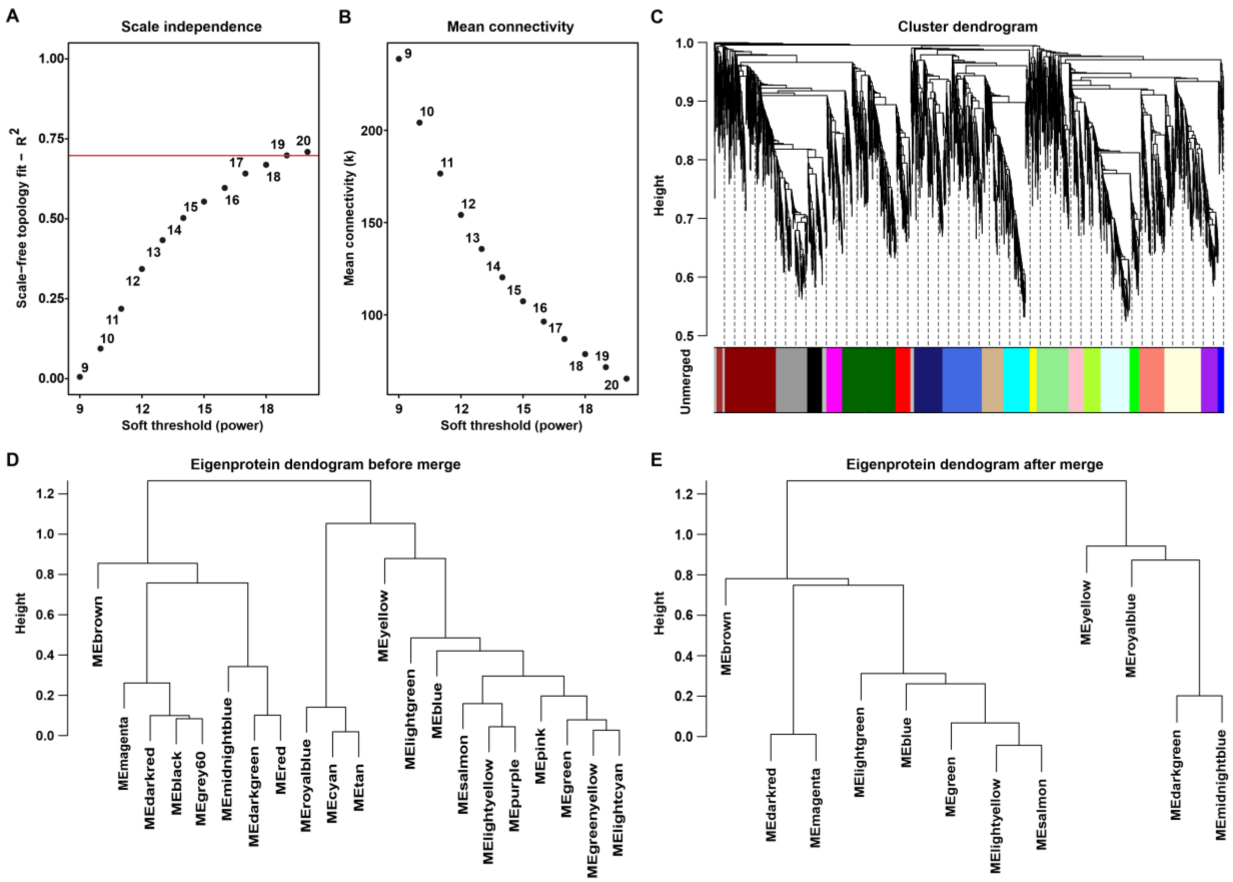


**Figure S3: Weighted protein co-expression network with scale-free topology generates co-expressed modules that can be summarized by eigenproteins. (A)** Network fit to a scale-free topology across soft power thresholds. Fit is determined by the log-log correlation between the connectivity probability P(k) and connectivity (k). The red line indicates the threshold selected (20). **(B)** Mean network connectivity (k) across soft power thresholds. **(C)** Cluster dendrogram of the network, where height represents the dissimilarity of clusters across modules. Each module is shown by a different color. **(D)** Simplified dendrogram from (C) containing 21 co-expressed modules, where each module is represented by its eigenprotein (ME). **(E)** Eigenprotein dendrogram after merging similar modules with a pearson correlation > 0.85, yielding 12 co-expressed modules.

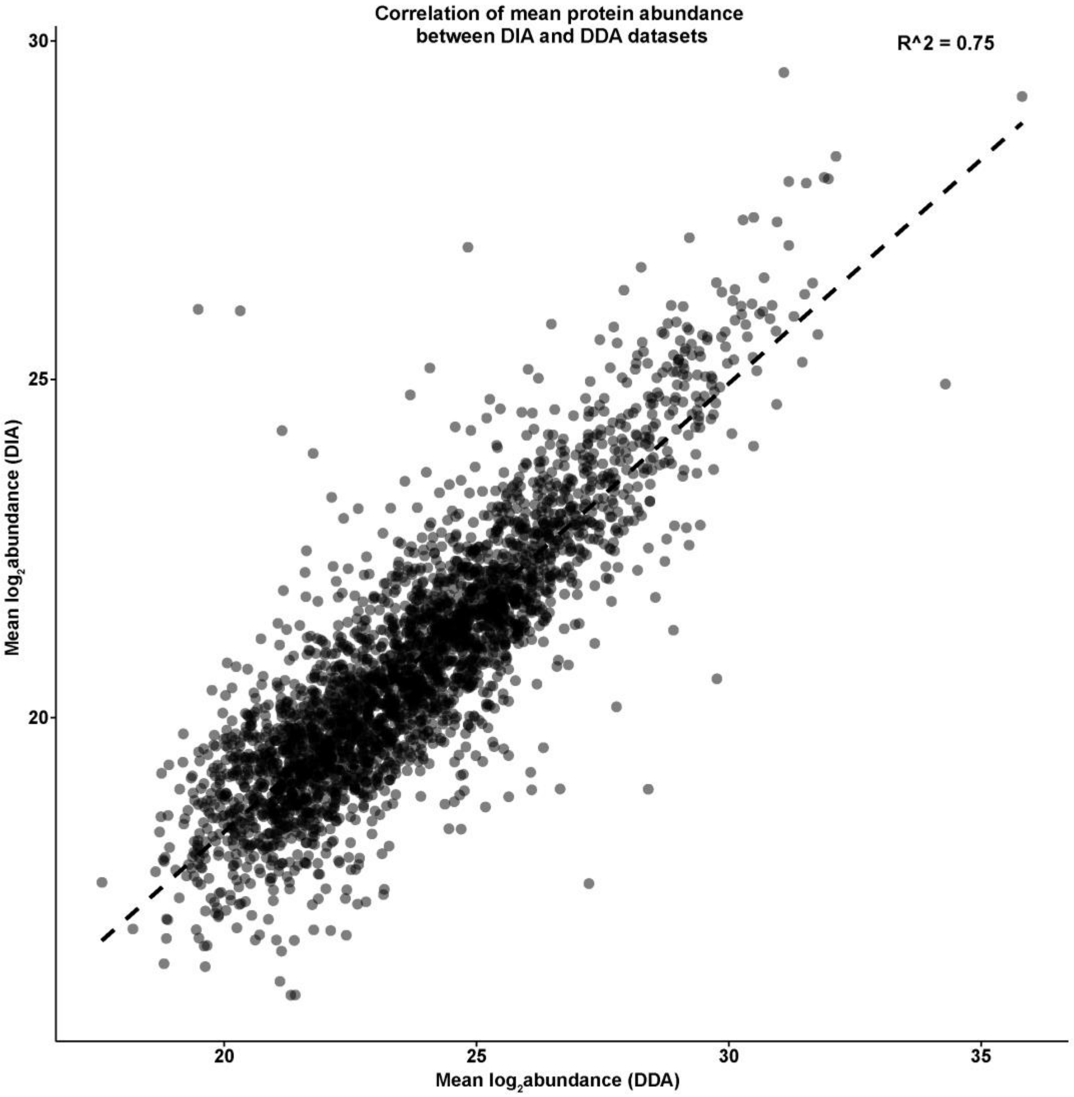


**Figure S4: Protein detection and abundance is similar across DIA and DDA protein acquisition methods.** Correlation of mean $\log_2$ protein abundance of the 3,027 proteins present in all samples across DIA and DDA datasets. The dashed line represents the line of best fit.

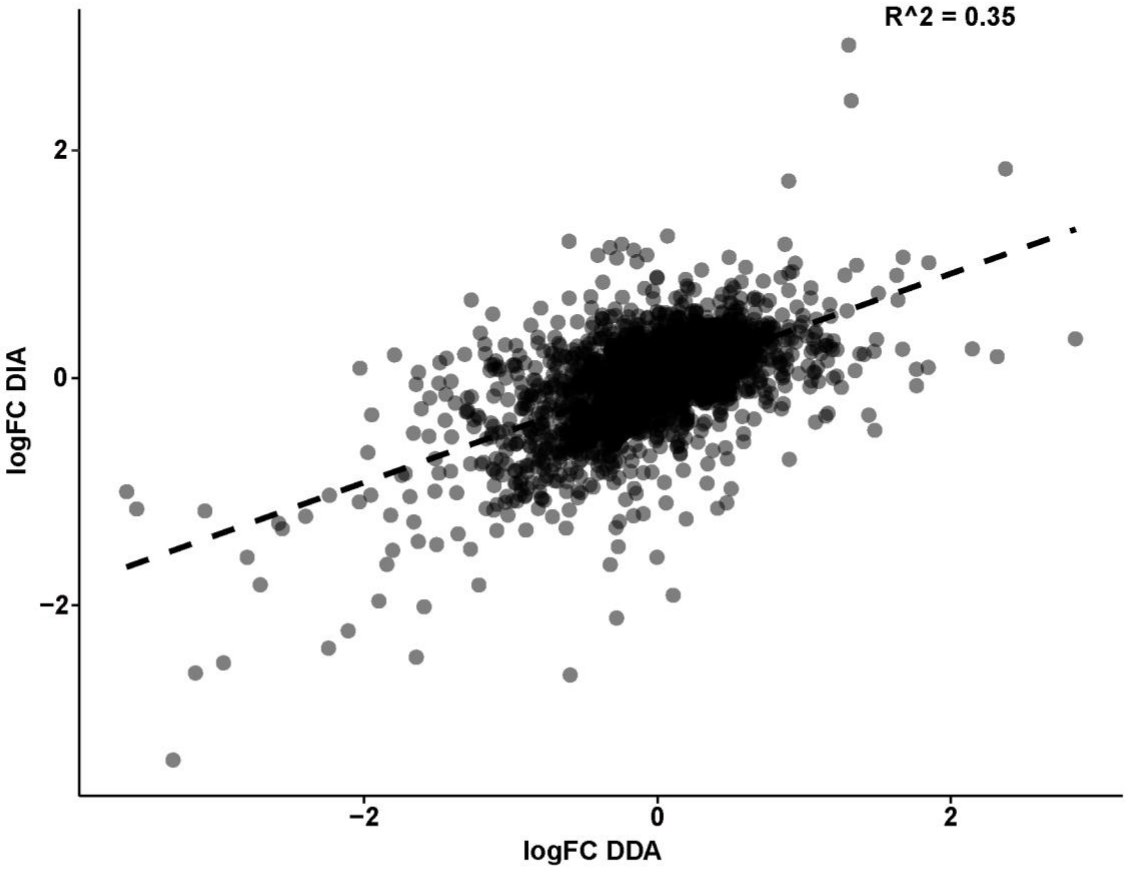


**Figure S5: Response to DOX between DDA and DIA protein acquisition methods is correlated.** $Log_2$ fold change between DOX and VEH for proteins detected and imputed using the Data-Dependent Acquisition (DDA) and Data-Independent Acquisition (DIA) methods is shown. The dashed line indicates the best fit line.

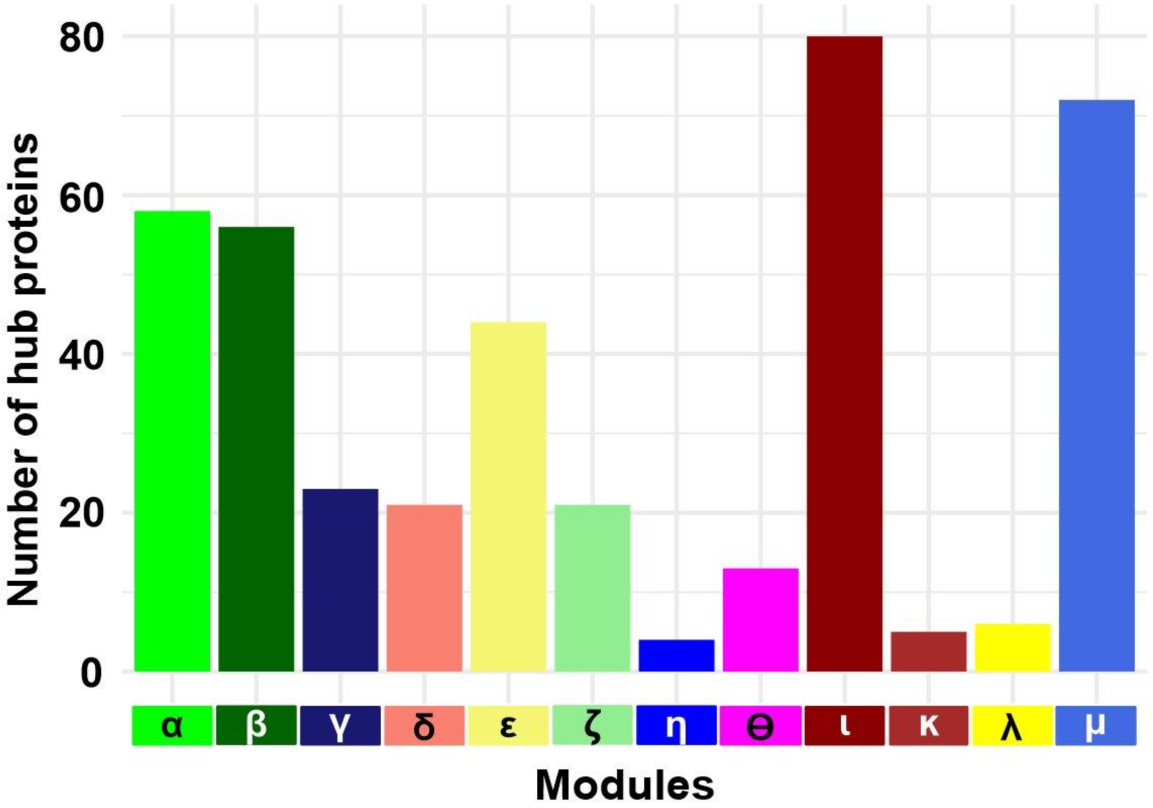


**Figure S6: Hub proteins are represented across all co-expressed modules.** Modules are ordered from α, the module with the strongest correlation to DOX, to μ, the module with the weakest correlation to DOX.

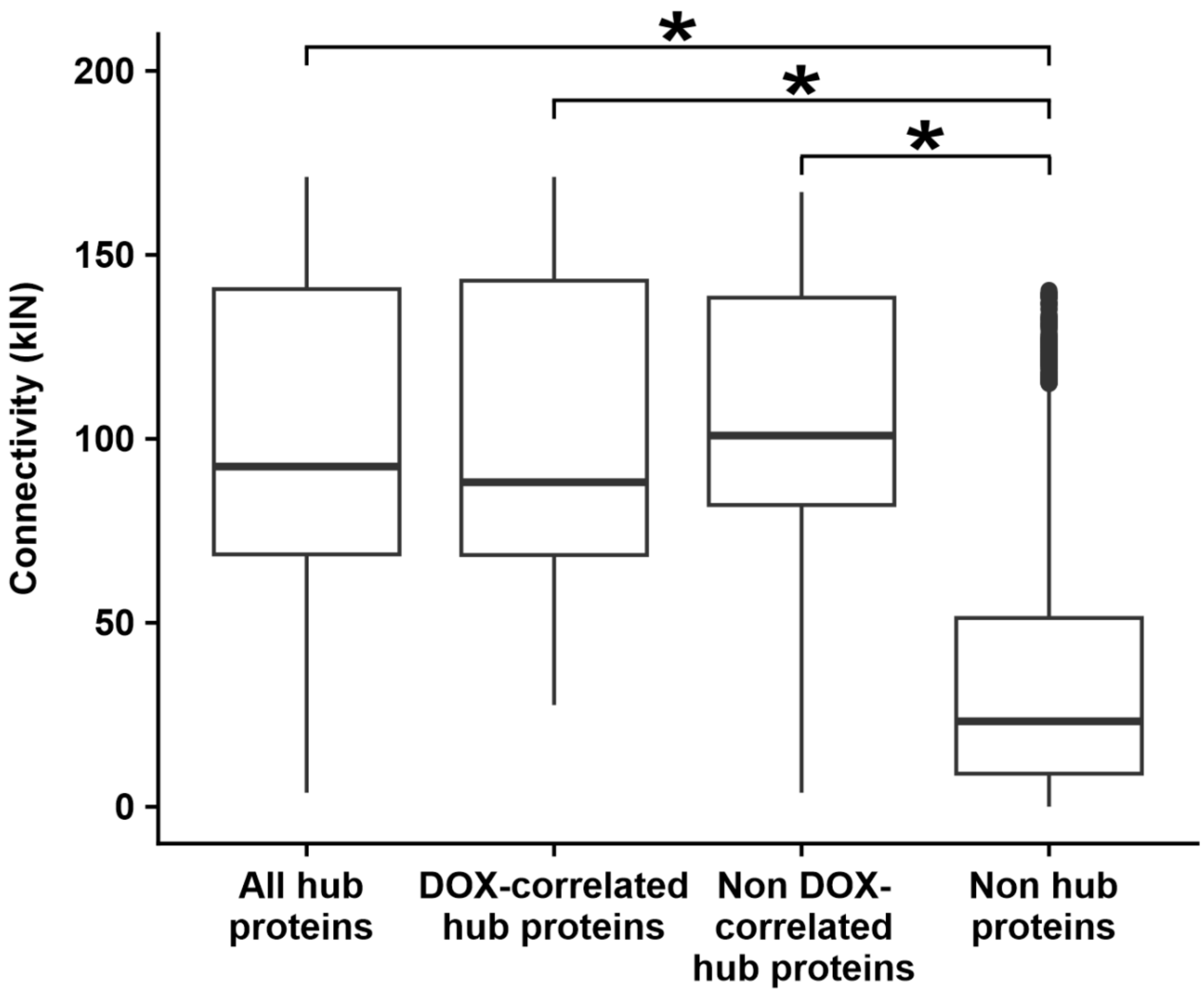


**Figure S7: Hub proteins have greater intramodular connectivity than non-hub proteins irrespective of DOX-correlation status.** Distribution of connectivity scores (kIN) for four categories of proteins: All hub proteins (n = 403), DOX-correlated hub proteins (n = 202), non-DOX-correlated hub proteins (n = 201), and network proteins that are non-hub proteins (n = 3,775). Asterisk denotes a statistically significant difference in kIN between conditions ($P < 0.05$).

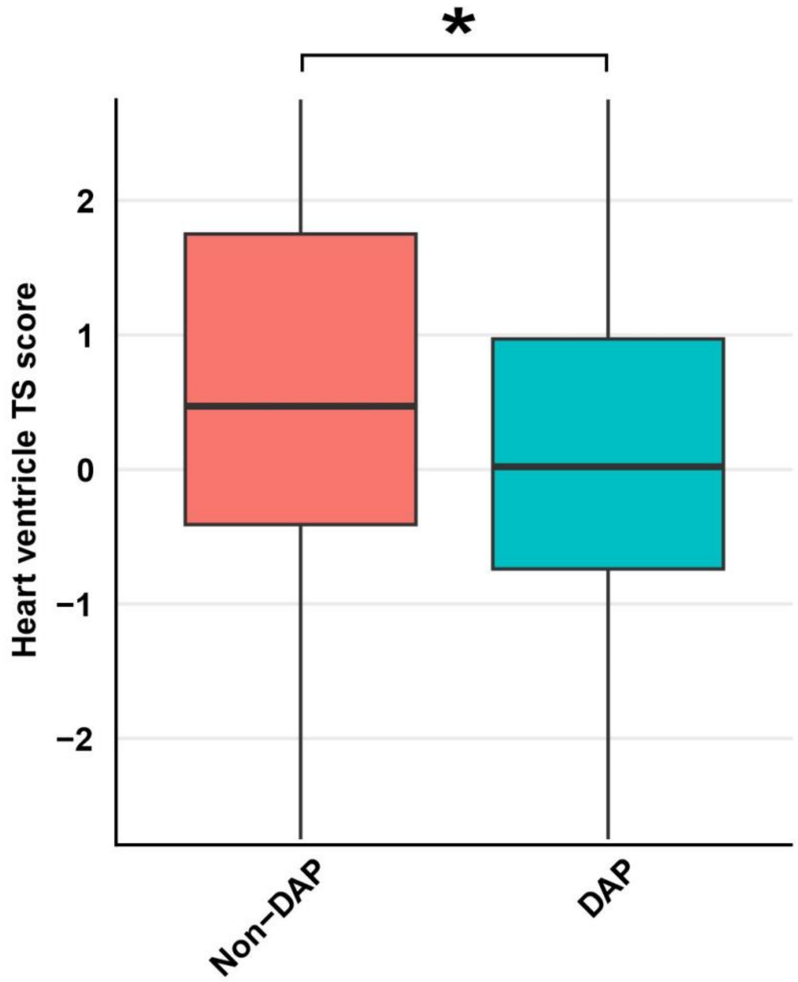


**Figure S8: Differentially abundant proteins are less specific to heart ventricle than proteins that are not differentially abundant.** Heart ventricle tissue-specificity (TS) scores for differentially abundant proteins (DAPs) and Non-DAPs (Uhlén *et al*., Science, 2015). Lower scores indicate that the protein is expressed across more tissues. Asterisk denotes a statistically significant difference in TS scores between groups ($P < 0.05$).

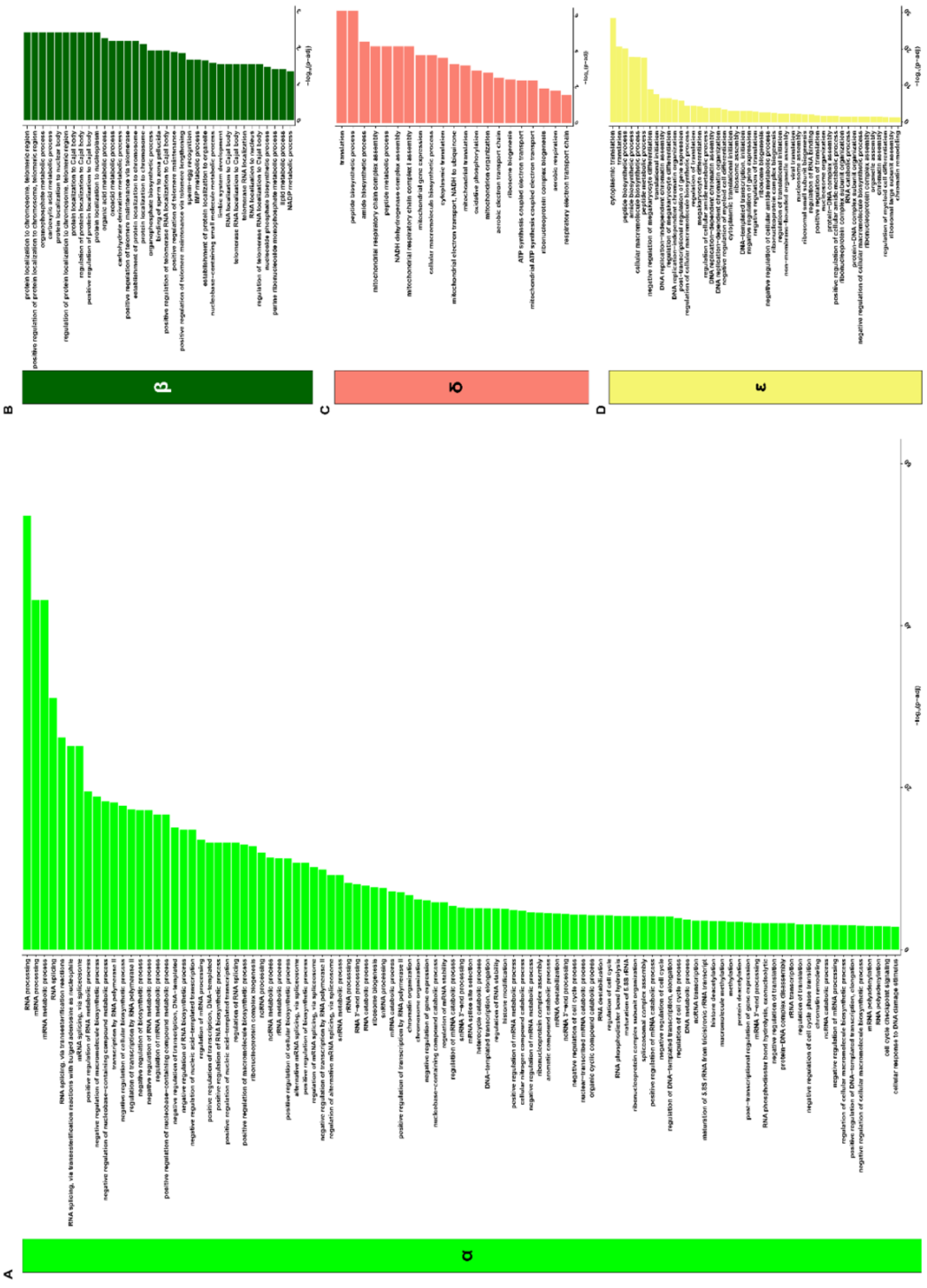

**Figure S9: DOX-correlated modules are enriched for many biological processes. (A)** Biological processes enriched in the α module (See Table 3 for complete term list). **(B)** Biological processes enriched in the β module. **(C)** Biological processes enriched in the δ module. **(D)** Biological processes enriched in the ε module. Enriched processes are defined by Fisher's exact test adjusted $P < 0.05$.

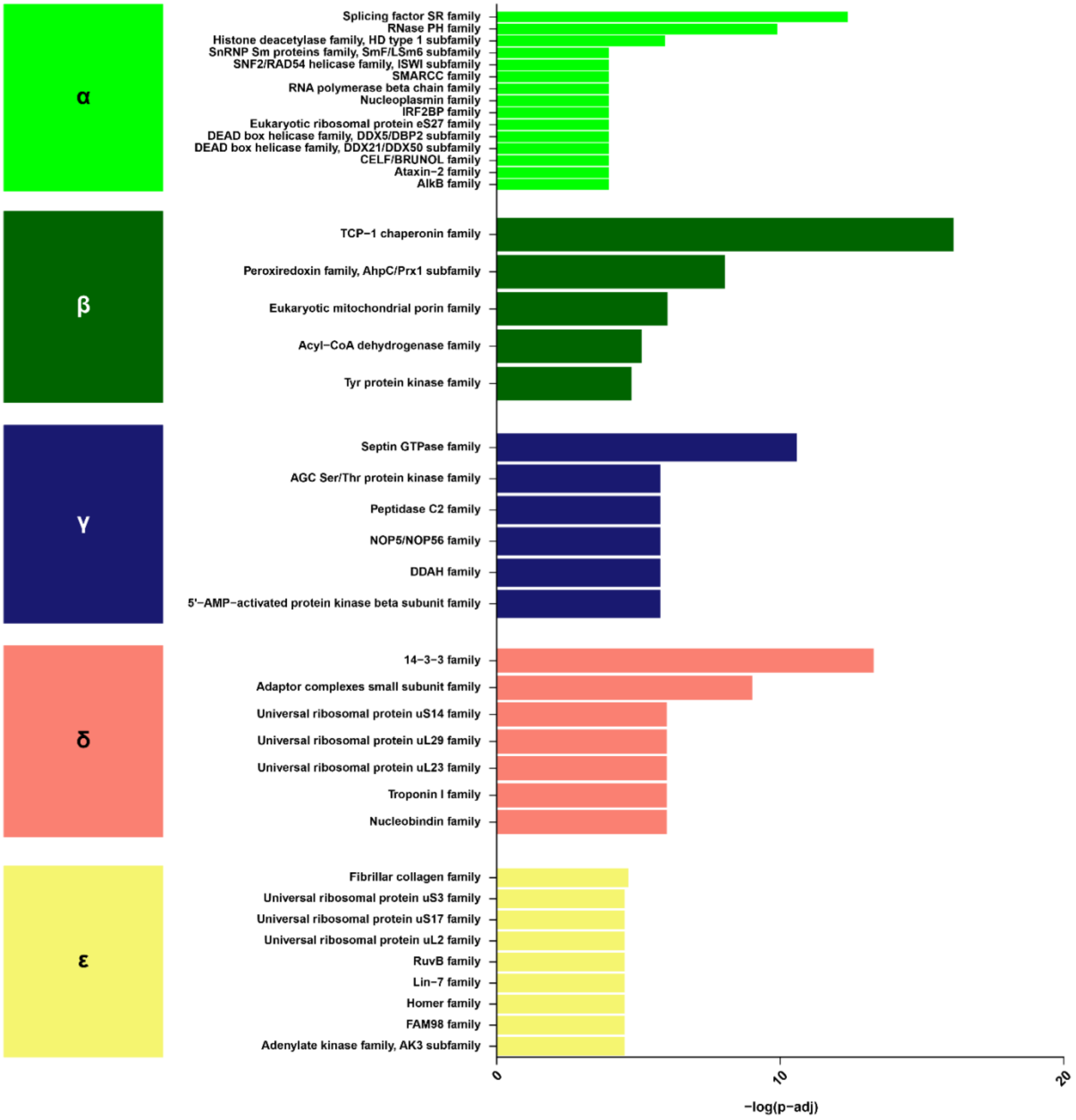


**Figure S10: DOX-correlated modules differ by their enriched protein families.** DOX-correlated modules α, β, γ, δ and ε are ordered by their correlation to DOX. Enrichment of protein families is determined by Fisher's exact test $P < 0.05$. The top five enriched families in each module were selected for visualization. Differences in the number of families depicted across modules are due to tied $P$ values from the Fisher's exact test.

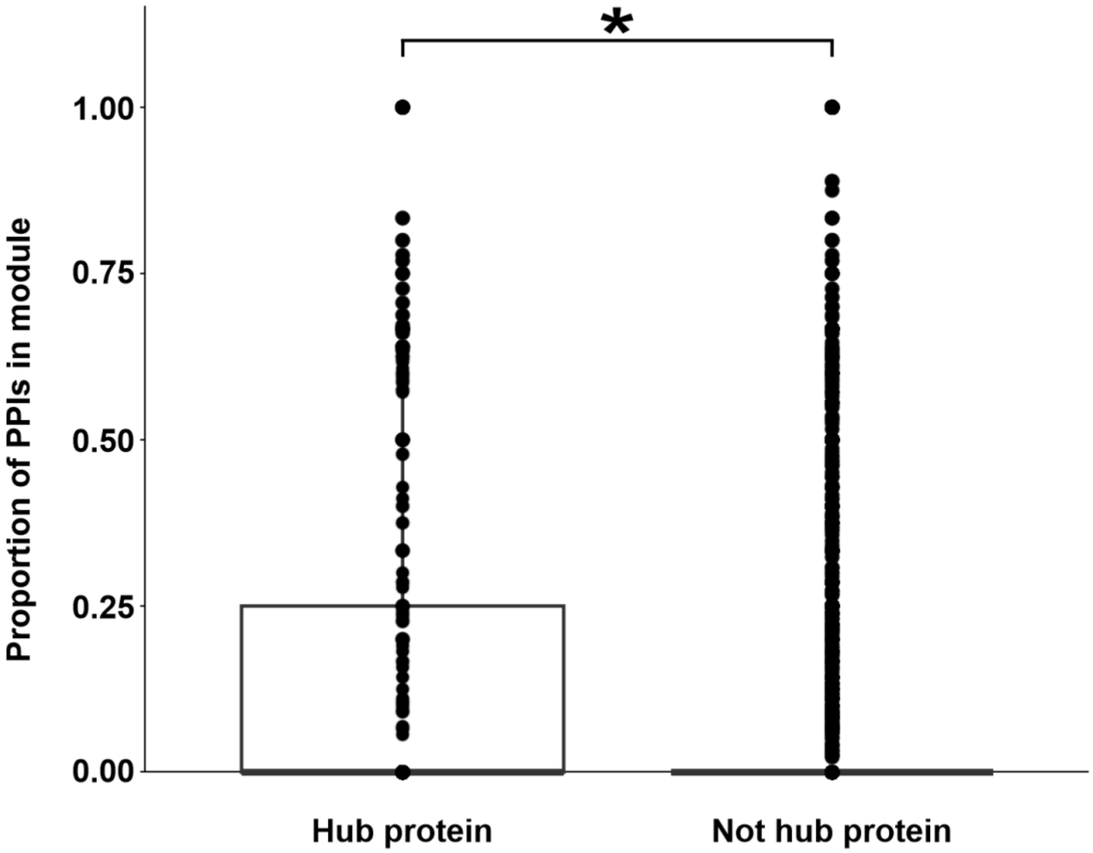


**Figure S11: Hub proteins are more likely to be co-expressed with proteins they physically interact than proteins that are not hubs.** Proportion of protein-protein interactions (PPIs) where both interactors are contained within the same co-expression module for hub proteins and non-hub proteins. PPIs of expressed proteins were obtained from STRINGdb (Szklarczy *et al.*, Nucleic acids research, 2023), where a confidence score of 0.9 ≥ is used as the threshold for interaction. Asterisk denotes a statistically significant difference in the proportion of PPIs within the same module between hub and non-hub proteins ($P < 0.05$).

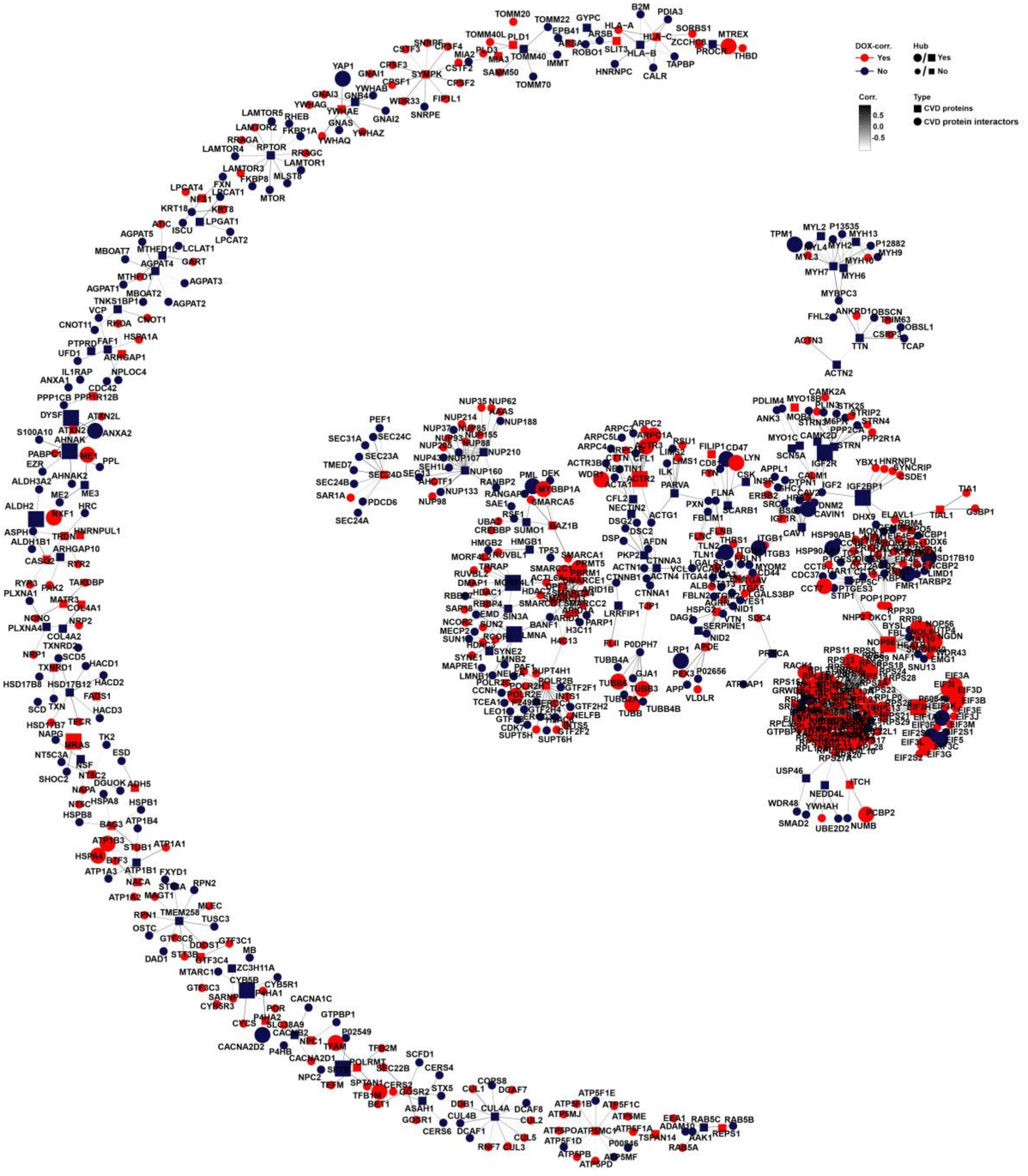


**Figure S12: Functionally annotated CVD-PPI network within the context of the DNA damage response.** Protein-protein interaction (PPI) network for CVD risk proteins (square) and CVD risk protein interactors (circle) expressed within the co-expression network. Edges represent the weighted correlation between interaction pairs. Node size indicates if a protein is a hub (large icon) or not a hub (small icon) protein. Color denotes if a protein is DOX-correlated (red) or not DOX-correlated (blue).
